# Shared Binding Site but Divergent Resistance Profiles Uncover Novel Resistance Mechanisms in *Plasmodium* HSP90 Inhibitors

**DOI:** 10.1101/2025.10.30.684657

**Authors:** Fu-Hsuan Ko, Amanda K. Lukens, Mariana Laureano de Souza, Alex Foy, Jason Hsiao, Tolla Ndiaye, John Okombo, Tomas Yeo, Heekuk Park, Anne-Catrin Uhlemann, Nonlawat Boonyalai, Krittikorn Kumpornsin, Gareth Girling, Charisse Pasaje, Luiz Godoy, Sabine Ottilie, Gregory L. Durst, Arnab Chatterjee, Jacquin Niles, Marcus Lee, David A. Fidock, Dyann F. Wirth, Elizabeth A. Winzeler

## Abstract

Drug resistance is a widespread problem across therapeutic areas including malaria, but what accounts for resistance propensity remains poorly understood. Here, we reveal that two HSP90 inhibitors targeting the identical ATP-binding site exhibit dramatically different resistance profiles in *P. falciparum*. Geldanamycin readily selected 10 distinct resistance mutations conferring up to 22-fold resistance, while AUY-922 required 44 weeks to yield a single A41S mutation with only 2-fold resistance to AUY-922 but not to geldanamycin. Resistance mapping in parasites and yeast revealed geldanamycin resistance mutations distributed throughout the binding pocket whereas AUY-922 resistance mutations localized close to the ATP-binding site. Unexpectedly, the A41S mutation enhanced AUY-922 binding affinity without changing geldanamycin binding. *In silico* analysis suggested this enhancement occurs through additional hydrogen bonding, yet stronger binding correlated with resistance. In yeast, A41S had opposite effects, hypersensitizing cells to all HSP90 inhibitors tested. Additionally, conditional HSP90 knockdown increased geldanamycin sensitivity but left AUY-922 activity unaffected, indicating different target dependencies despite shared binding sites. Based on these data, we propose a multi-target hypothesis where AUY-922’s lower resistance risk stems from engaging multiple HSP90 family members. Our findings reveal how enhanced drug-target binding can paradoxically correlate with resistance and demonstrate that resistance risk cannot be predicted from binding site identity alone, providing insights for developing more durable drugs across therapeutic areas.

## Introduction

Drug resistance threatens therapeutic efficacy across multiple medical fields, with malaria representing a critical example where resistance evolution undermines treatment success. *Plasmodium falciparum* causes the most severe malaria cases, and despite extensive control efforts, the disease continues to affect millions annually with hundreds of thousands of deaths (Venkatesan 2025). Resistance has emerged against all deployed antimalarial drugs within years to decades of introduction, creating cycles of drug development and resistance that threaten elimination goals (Haldar et al. 2018; Rosenthal et al. 2024). This creates an urgent need for new therapeutic approaches that can overcome existing resistance mechanisms and ideally exhibit enhanced durability against future resistance evolution (Siqueira-Neto et al. 2023). Current antimalarial drug discovery efforts emphasize the need to address fundamental questions in drug development: can resistance risk be predicted from target characteristics and do certain design approaches yield compounds with inherently higher barriers to resistance evolution? Understanding these principles has implications beyond malaria for any therapeutic area where drug resistance threatens treatment durability.

Heat shock protein 90 (HSP90) has attracted attention as a potential antimalarial target due to its essential role as a molecular chaperone. As a highly conserved ATPase, HSP90 is involved in the folding, maturation, and maintenance of numerous client proteins critical for cellular function, making it an attractive target for therapeutic intervention (Roy et al. 2012; Ramdhave et al. 2013). HSP90 protein plays crucial roles in protein quality control, signal transduction, and stress response pathways that are essential for parasite survival and development. *Plasmodium* parasites express multiple essential HSP90 isoforms in different organelles, including the ER-resident GRP94 (glucose-regulated protein 94), which may offer additional therapeutic opportunities for targeting. Several HSP90 inhibitors have demonstrated potent antimalarial activity against *P. falciparum* with half maximal inhibitory concentration (IC_50_) values often in the nanomolar range (Murillo-Solano et al. 2017; Posfai et al. 2018). Some of these compounds were identified from the Calibr REFRAME collection, a library of small molecules that have reached clinical development or undergone significant preclinical profiling (Janes et al. 2018). These include AUY922 (luminespib), which had previously advanced to human clinical trials for cancer treatment. However, the precise mechanisms by which these inhibitors achieve their antimalarial effects and their susceptibility to resistance development remain incompletely characterized.

Most HSP90 inhibitors target the highly conserved ATP-binding pocket, leading to the expectation that compounds binding to this site would exhibit similar resistance profiles. The ATP-binding domain is structurally conserved across species (Southworth and Agard 2008) and represents the most commonly targeted site for HSP90 inhibition (Li and Luo 2023). While this conservation might suggest similar resistance mechanisms across species, resistance development can be more complex. Resistance can arise through multiple mechanisms beyond target binding, including altered drug kinetics, cellular distribution, and protein function. Understanding how these factors influence resistance development is crucial for predicting the useful therapeutic lifespan of HSP90-targeted antimalarials and for designing compounds with improved resistance profiles.

We employed comprehensive resistance profiling including *in vitro* resistance evolution experiments, minimum inoculum for resistance assays, CRISPR-Cas9 gene editing, conditional knockdown studies, and biophysical binding analysis. Surprisingly, we found that despite targeting the same binding site, these compounds exhibited dramatically different resistance profiles, with AUY-922 showing an exceptionally high barrier to resistance development compared to geldanamycin. We identify a paradoxical resistance mechanism where enhanced drug binding correlates with reduced antimalarial efficacy, challenging conventional structure-activity relationships. These findings demonstrate that resistance risk cannot be predicted solely from target binding site identity and reveal how compounds with similar binding modes can exhibit different evolutionary vulnerabilities, with implications for designing more durable antimalarial therapies.

## Results

### Evaluation of Two Structurally Distinct HSP90 Inhibitors

We first evaluated two structurally distinct HSP90 inhibitors that bind to the ATP-binding pocket against asexual blood-stage *P. falciparum* Dd2 parasites (Figure 1A). AUY-922 (luminespib) is a resorcinol-based inhibitor that was originally developed for cancer treatment and reached Phase II clinical trials (Felip et al. 2018), while geldanamycin is an ansamycin antibiotic natural product that has served as a prototype HSP90 inhibitor. Despite targeting the same binding site, AUY-922 showed superior potency (IC₅₀ = 20 nM) compared to geldanamycin (IC₅₀ = 158 nM), an ∼8-fold difference in activity (Figure 1B). To assess their potential as multi-stage antimalarials, we evaluated activity against liver-stage parasites using a *P. berghei* luciferase reporter system (Swann et al. 2016). AUY-922 maintained its potency advantage in liver stages (IC₅₀ = 111 nM) compared to geldanamycin (IC₅₀ = 729 nM) (Figure 1C). Neither compound exhibited significant cytotoxicity against human HepG2 hepatocytes at concentrations up to 10 µM, providing initial selectivity data for these HSP90 inhibitors in the hepatocyte model system (Figure 1D). We also characterized radicicol, a resorcylic acid lactone natural product that targets the same ATP-binding pocket as AUY-922 and geldanamycin. Radicicol demonstrated antimalarial activity against blood-stage *P. falciparum* (IC₅₀ = 1.91 µM) and liver-stage *P. berghei* parasites (IC₅₀ = 13.92 µM), with cytotoxicity against human hepatocytes at higher concentrations (IC₅₀ = 8.05 µM) (Figure S1). This third HSP90 inhibitor provided additional validation of the target and served as a useful comparator in subsequent resistance and conditional knockdown experiments.

**Figure 1.**
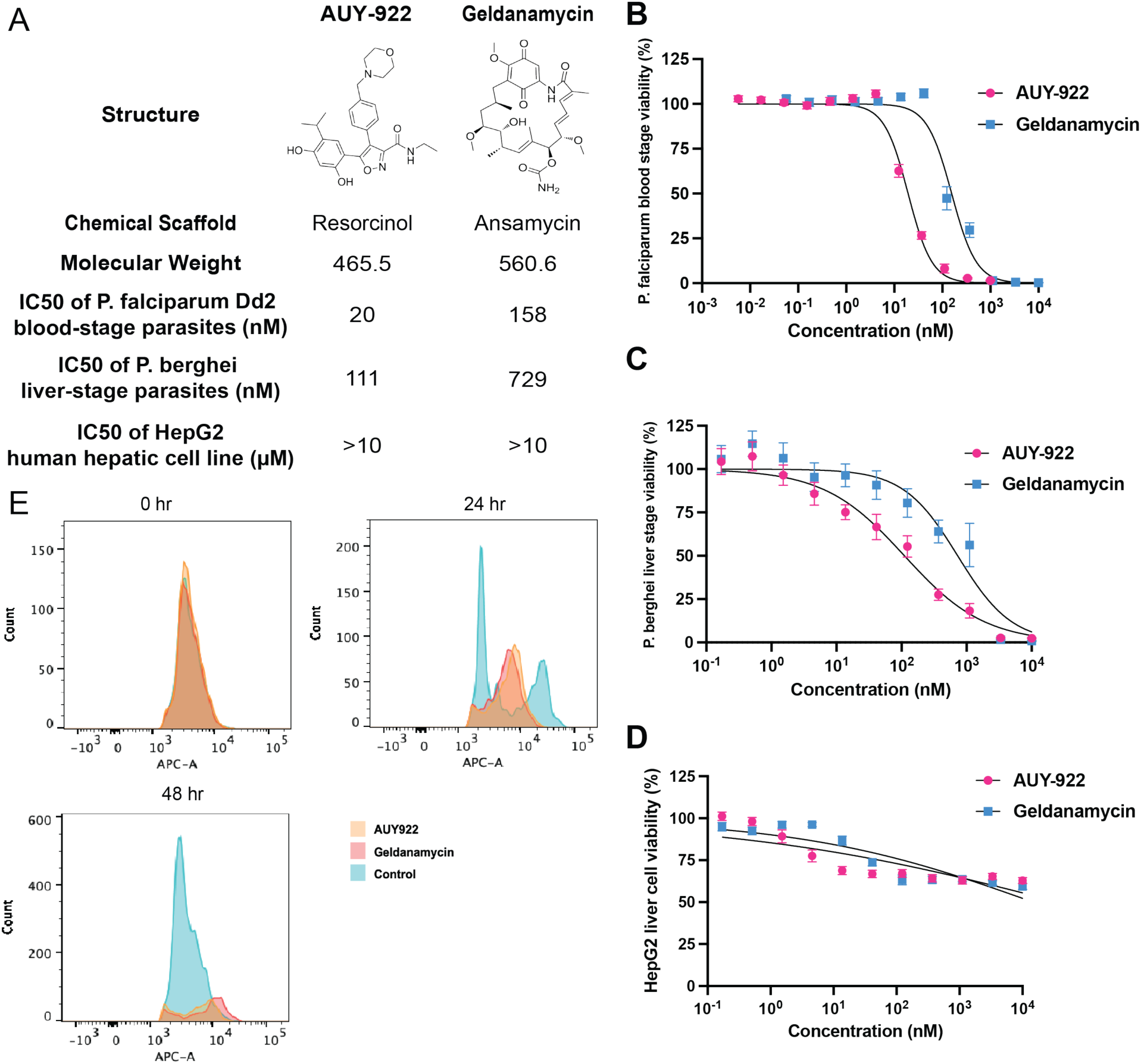
Characterization and mechanism of action studies of HSP90 inhibitors. (A) Chemical structures and biological activity profiles of two structurally diverse HSP90 inhibitors: AUY-922 (resorcinol-based) and geldanamycin (ansamycin), summarizing blood stage, liver stage, and cytotoxicity data from panels B-D. (B) Dose-response curves for AUY-922 and geldanamycin against *P. falciparum* Dd2 strain. IC_50_ values were determined using a 72-hour SYBR Green-based growth assay. (C) Life stage-specific activity assessment showing IC₅₀ values of AUY-922 and geldanamycin against liver stage *P. berghei* parasites in a luciferase-based assay. (D) Cytotoxicity assessment in HepG2 human hepatocytes showing that both AUY-922 and geldanamycin exhibited minimal toxicity, with no measurable IC₅₀ values obtained at concentrations up to 10 µM. (E) Flow cytometry analysis using SYTO61 DNA staining showing intraerythrocytic developmental stages of *P. falciparum* after drug treatment. Histograms demonstrate parasite progression through the blood stage lifecycle, with distinct profiles for ring, trophozoite, and schizont stages.

To elucidate the developmental stage specificity of HSP90 inhibitor action, we monitored parasite progression through the intraerythrocytic cycle using flow cytometry with SYTO61 DNA staining (Fu et al. 2010). Tightly synchronized ring-stage parasites were treated with 5× IC₅₀ concentrations of either AUY-922 or geldanamycin, and DNA content was assessed at 24 and 48-hours post-treatment (Figure 1E). At 24 hours, control parasites exhibited the characteristic dual-peak pattern reflecting normal trophozoite development and nuclear division. In stark contrast, both AUY-922- and geldanamycin-treated parasites remained predominantly arrested in ring stage, displaying reduced signal intensity indicative of developmental arrest and partial parasite death. By 48 hours, control parasites had completed their developmental cycle and reinvaded fresh erythrocytes, showing the expected single peak characteristic of new ring-stage infections. Drug-treated populations, however, displayed diminished and scattered signal patterns, suggesting widespread parasite death following prolonged developmental arrest. The similar developmental arrest profiles induced by both compounds suggest they interfere with critical cellular processes required for the ring-to-trophozoite transition, consistent with disruption of essential chaperone functions during this metabolically active developmental period. This stage-specific vulnerability aligns with the known importance of protein folding and quality control mechanisms during the rapid parasite growth and metabolic reorganization that characterizes trophozoite development.

### Target Validation by in vitro Resistance Evolution in *P. falciparum*

In vitro evolution experiments provide a powerful approach to investigate mechanisms of drug resistance and potential drug targets in an organismal context. When parasites acquire resistance-conferring mutations in a suspected target after compound exposure, it can provide valuable evidence supporting on-target activity, though resistance mutations can also arise in other pathways that affect drug efficacy. Previous evolution experiments from our lab showed that *S. cerevisiae* readily acquired multiple resistance mutations to AUY-922, all located in the N-terminal ATP-binding domain of ScHSP90 (Table 1) (Ottilie et al. 2022). Given the high conservation of this domain between species, we anticipated finding similar resistance patterns in *P. falciparum*. However, the malaria parasite exhibited different behavior. Despite subjecting *P. falciparum* to an exceptionally prolonged selection period of 44 weeks with gradually increasing AUY-922 concentrations, resistance development proved remarkably difficult. This extensive evolution experiment yielded only a single resistant clone showing a 4-fold increase in IC₅₀ (Figure 2A). Whole-genome sequencing identified an A41S amino acid change in the ATP-binding domain of PfHSP90, along with additional mutations detected elsewhere in the genome. The nearly year-long selection pressure required to obtain even a single resistant clone in *P. falciparum* indicates an exceptionally high barrier to AUY-922 resistance in the malaria parasite compared to yeast.

**Figure 2.**
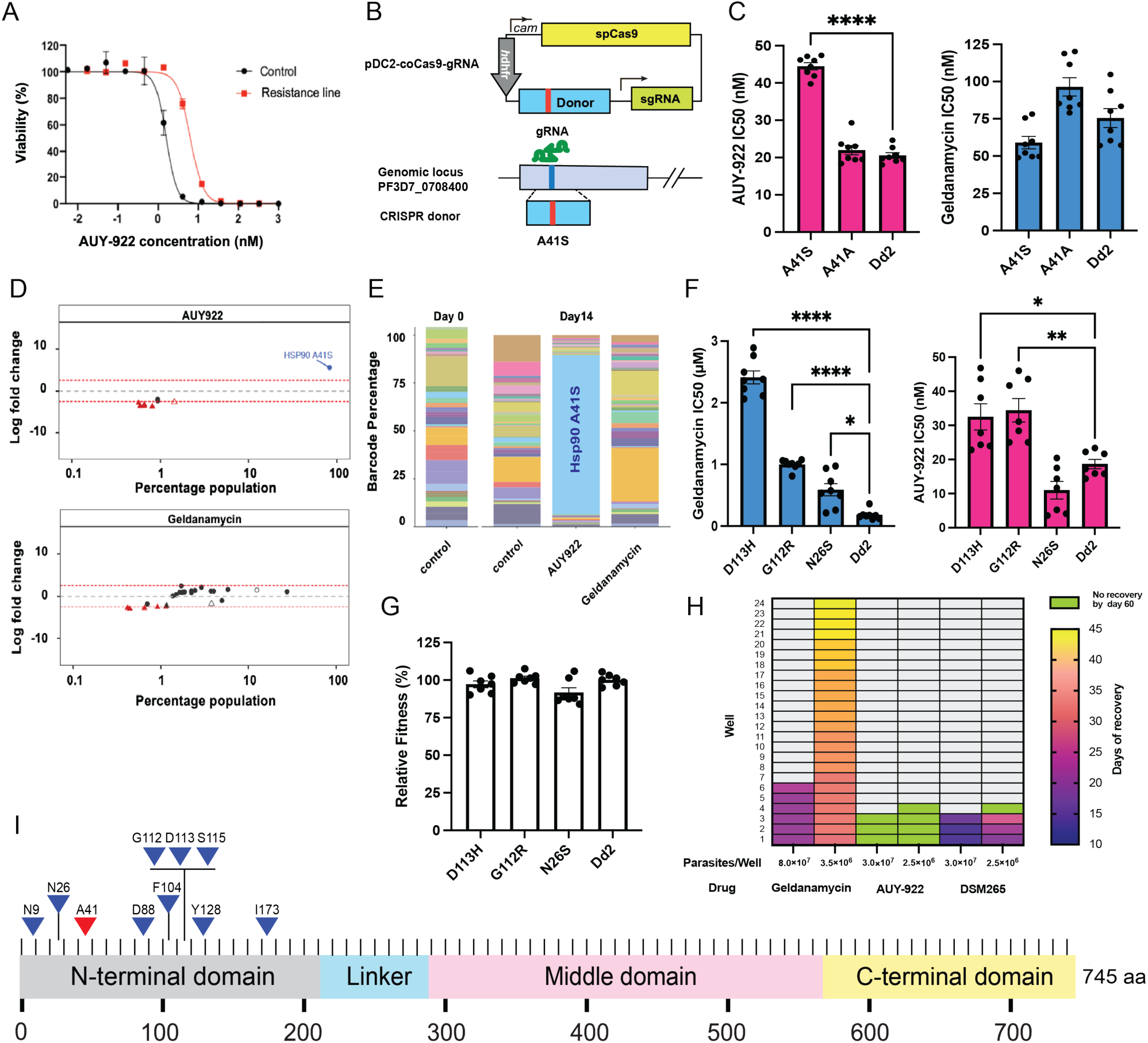
Resistance analysis and validation of HSP90 inhibitors. **(A)** Dose-response curves of parental Dd2 and AUY-922 resistant line evolved over 44 weeks. **(B)** CRISPR-Cas9 strategy for generating the A41S mutation in the PfHSP90 genomic locus. **(C)** IC₅₀ values of CRISPR-engineered parasites against AUY-922 (left) and geldanamycin (right). **(D)** AReBar competitive growth assay showing log₂ fold-change analysis under AUY-922 (top) and geldanamycin (bottom) selection pressure. Each dot (circles: Dd2 background, triangles: 3D7 background) represents a barcoded mutant (filled) or wild type (empty) line, with parasites <0.1% not shown (see Table S3). **(E)** Barcode percentage composition at day 0 and day 14 after drug selection. Each bar represents the proportion of a barcoded line. **(F)** IC₅₀ values of geldanamycin-resistant clones against geldanamycin (left) and AUY-922 (right). **(G)** Relative fitness assessment of geldanamycin-resistant clones. **(H)** Minimum Inoculum for Resistance (MIR) assay results displayed as a heatmap. Color gradient indicates days of recovery (10-45 days), lime indicates no recovery by day 60, and light grey indicates no data point. **(I)** Schematic representation of PfHSP90 domain organization showing resistance mutation distribution. AUY-922 resistance mutation A41S (red triangle) and geldanamycin resistance mutations (blue triangles).

**Table 1.** Mutations identified in in vitro evolution or MIR experiments and IC_50_ fold-shift with AUY-922 and geldanamycin in *P. falciparum* and *S. cerevisiae*. Yeast mutations (*S. cerevisiae* ABC16-Monster) are indicated with asterisk. IC_50_ fold-shifts represent resistance of mutant strains compared to wild-type Dd2 (*P. falciparum*) and wild-type ABC16-Monster (*S. cerevisiae*).

| AUY-922 |  |  | Geldanamycin |  |  |
| --- | --- | --- | --- | --- | --- |
| Nucleotide | Amino Acid | IC<sub>50</sub> shift- fold | Nucleotide | Amino Acid | IC<sub>50</sub> shift -fold |
| aaC/aaA | A41S | 4.0 | aaC/aaA | N9K | 22.3 |
| *aCc/aAc | *T171N | *6.7 | Gat/Tat | D88Y | 3.8 |
| *Gcc/Ccc | *A112P | *6.1 | GgA/TgG | G112W | 1.8 |
| *gaT/gaG | *D79G | *37.8 | aAc/aGc | N26S | 3.1 |
|  |  |  | Gga/Cga | G112R | 5.9 |
|  |  |  | Gat/Cat | D113H | 12.2 |
|  |  |  | tTt/tAt | F104Y | 5.0 |
|  |  |  | tCt/tAt | S115Y | 3.3 |
|  |  |  | Tat/Cat | Y128H | 2.6 |
|  |  |  | Att/Ttt | I173F | 5.3 |

Because the A41S *P. falciparum* mutation was a low-probability singleton mutation and other mutations were present (Table 1), we introduced this mutation, or alternatively a silent A41A control mutation into a clean genetic background using CRISPR Cas9 engineering (Figure 2B). The edited parasites were validated and cloned and then tested for resistance to AUY-922. These data showed the A41S but not the control silent mutation was indeed able to confer statistically significant, 2x shift in IC_50_ to AUY-922 (Figure 2C). Notably, the A41S mutation did not confer cross-resistance to geldanamycin, suggesting distinct resistance mechanisms for these two HSP90 inhibitors.

To further validate the specificity of AUY-922 for HSP90 and test whether the compound also hits other known antimalarial targets, we employed the Antimalarial Resistome Barcode (AReBar) platform, which allows simultaneous tracking of a diverse panel of resistance mutations in a mixed population (Carrasquilla et al. 2022). In this system, the pool of parasite lines encompasses >30 different modes of action, with various validated resistance mutations to different antimalarial compounds, including mutations conferring resistance to artemisinin, atovaquone, and other clinical, preclinical and experimental antimalarials, including with our HSP90 A41S mutant (Table S3). Barcode frequency analysis of mixed populations under drug pressure (3×IC_50_ for 14 days) revealed highly specific drug-resistance interactions (Figure 2D). The differential analysis established HSP90 A41S as the only variant showing significant enrichment (>2.5 log₂ fold change) in AUY-922-treated cultures, corresponding to parasites harboring the HSP90 A41S mutation reaching over 90% of the population (Figure 2E), whereas all other resistance mutations were not appreciably enriched. In contrast, geldanamycin-treated populations showed no significant enrichment of the A41S variant, further confirming the lack of cross-resistance between these compounds (Fig. 2D, E).

We also examined geldanamycin resistance in malaria parasites using an early cloning protocol in which parasites are exposed to geldanamycin and then rapidly isolated before a clone would have an opportunity to grow out and outcompete everything else (Lim et al. 2016). In contrast to AUY-922, multiple geldanamycin-resistant parasites were readily obtained, with clones all showing increased IC_50_ values. Targeted HSP90 sequencing of six of these clones confirmed that all had acquired mutations conferring resistance to geldanamycin in the *P. falciparum* HSP90 gene. The identified mutations, found in separate clones, included N9K, D88Y, G112W, N26S, G112R, and D113H (Table 1). These resistant clones exhibited increased IC_50_ values for geldanamycin, with fold changes ranging from 1.8 to 22.3 compared to the wild-type Dd2 parasite. A key finding is the D88Y mutation, which occurs at a position analogous to E88 in *S. cerevisiae* Hsp82, previously identified as a resistance site for geldanamycin derivatives (Millson et al. 2011). In both organisms, mutations at position 88 replace an acidic residue (aspartic acid in PfHSP90, glutamic acid in yeast) with a non-acidic one, suggesting a conserved mechanism of resistance through altered drug-protein interactions in the ATP binding pocket.

### Cross-resistance Testing Reveals Different Mechanisms of Action for HSP90 Inhibitors

To further characterize the resistance mechanisms, we conducted cross-resistance profiling. These geldanamycin-resistant clones showed minimal cross-resistance to both AUY-922 (Figure 2F) and radicicol (Figure S2A). The log2 fold-change analysis revealed distinct resistance patterns: while all three tested mutations (D113H, G112R, and N26S) showed strong resistance to geldanamycin (positive log2 fold changes), they exhibited minimal or no resistance to AUY-922 and radicicol, with some mutations even showing slight sensitization (negative log2 fold changes) to these compounds (Figure S2B). To validate the specificity of these resistance patterns, we tested the geldanamycin-resistant clones against artemisinin and GNF179, antimalarial compounds with distinct mechanisms of action (Bridgford et al. 2018; LaMonte et al. 2020). As expected, these clones maintained wild-type sensitivity to both control compounds (Figure S2C, S2D), confirming that the resistance was specific to geldanamycin and did not reflect general drug resistance mechanisms. Furthermore, we assessed the fitness costs of these resistance mutations by comparing the growth rates of geldanamycin-resistant clones to wild-type parasites under normal culture conditions. No significant differences in growth rates were observed (Figure 2G), indicating that the geldanamycin resistance mutations do not impose a substantial fitness cost under these conditions.

### Comparative MIR Analysis Reveals Differential Resistance Barriers

The differences between the evolution time and number of recovered resistant clones for geldanamycin and AUY-922 were surprising given that both compounds should target HSP90. To further quantify the propensity for resistance development, we conducted Minimum Inoculum for Resistance (MIR) assays for both AUY-922 and geldanamycin (Figure 2H). This approach provides a quantitative measure of the likelihood of resistance development for each compound (Duffey et al. 2021). For AUY-922, no resistant parasites were observed in any of the wells throughout the 60-day period. This result indicates an MIR value of at least 10^8^, supporting a high barrier to resistance for AUY-922. In contrast, geldanamycin-treated cultures exhibited resistant parasites between days 21 and 45, with a minimum inoculum of 3.5 × 10^6^ parasites required for the development of resistance. This lower MIR value indicates a higher propensity for resistance development compared to AUY-922, consistent with our earlier observations from the in vitro evolution experiments. The control compound, DSM265, showed resistant parasites between days 13 and 27, corresponding to an MIR value of ∼3 × 10^6^. This result is consistent with its known propensity for resistance development (Phillips et al. 2015). Sequencing analysis of the resistant parasites from the geldanamycin MIR assay revealed four additional mutations in PfHSP90: F104Y, S115Y, Y128H, and I173F (Table 1). These new mutations, combined with those identified in our earlier experiments, highlight the diverse ways in which *P. falciparum* can develop resistance to geldanamycin, all involving modifications to PfHSP90.

Our evolution experiments yielded multiple resistance mutations across both compounds, and remarkably, all resistance-conferring mutations mapped exclusively to the N-terminal ATP-binding domain of PfHSP90 (Figure 2I). These resistance mutations include the single AUY-922 resistance mutation (A41S, shown in red) and multiple geldanamycin resistance mutations (N9K, D88Y, G112W, N26S, G112R, D113H, F104Y, S115Y, Y128H, and I173F, shown in blue). All mutations are located within the N-terminal ATP-binding domain near the expected binding site for geldanamycin based on crystal structures from other species (Whitesell et al. 2019; Prodromou et al. 1997). The exclusive localization of all resistance mutations within this conserved domain strongly supports HSP90 as the primary target for both compounds in *P. falciparum* and confirms that both inhibitors function through direct competition with ATP binding, consistent with their known mechanism of action in other organisms.

### Yeast Model Confirms HSP90 as Target for Geldanamycin and AUY-922

To further investigate the conservation of these resistance mutations, we used a yeast model. We systematically examined the functional impact of key mutations using CRISPR-based gene editing in yeast HSP90 (HSC82). After optimizing CRISPR/Cas9 vectors for the ABC16-Monster strain, which lacks 16 known adenosine triphosphate-binding cassette (ABC) transporters associated with drug resistance (Suzuki, St Onge, et al. 2011), we generated four mutations: A41S (the *P. falciparum* AUY-922 resistance mutation), E88Y (corresponding to the *P. falciparum* D88Y geldanamycin resistance mutation), A122P (identified in yeast AUY-922 evolution and equivalent to *P. falciparum* G112R geldanamycin resistance), and T171N (from yeast AUY-922 evolution).

The engineered yeast strains revealed definitive and striking differences in their drug responses. Testing against AUY-922 (Figure 3A) showed that the A41S mutation produced opposite effects between species—while it confers AUY-922 resistance in *P. falciparum*, it dramatically increased sensitivity in yeast. The A122P and T171N mutations both conferred strong resistance to AUY-922, while E88Y showed increased sensitivity. Against geldanamycin (Figure 3B), E88Y demonstrated clear resistance, while A122P and T171N also showed resistance. The A41S mutation again showed increased sensitivity to geldanamycin in yeast. Cross-resistance testing with radicicol revealed similar patterns (Figure S3A), with A122P and T171N conferring resistance while A41S and E88Y showed sensitization. The comprehensive fold-change analysis (Figure S3B) confirmed these distinct response patterns across all three HSP90 inhibitors, clearly demonstrating fundamental differences in HSP90 inhibitor mechanisms between yeast and *P. falciparum*.

**Figure 3.**
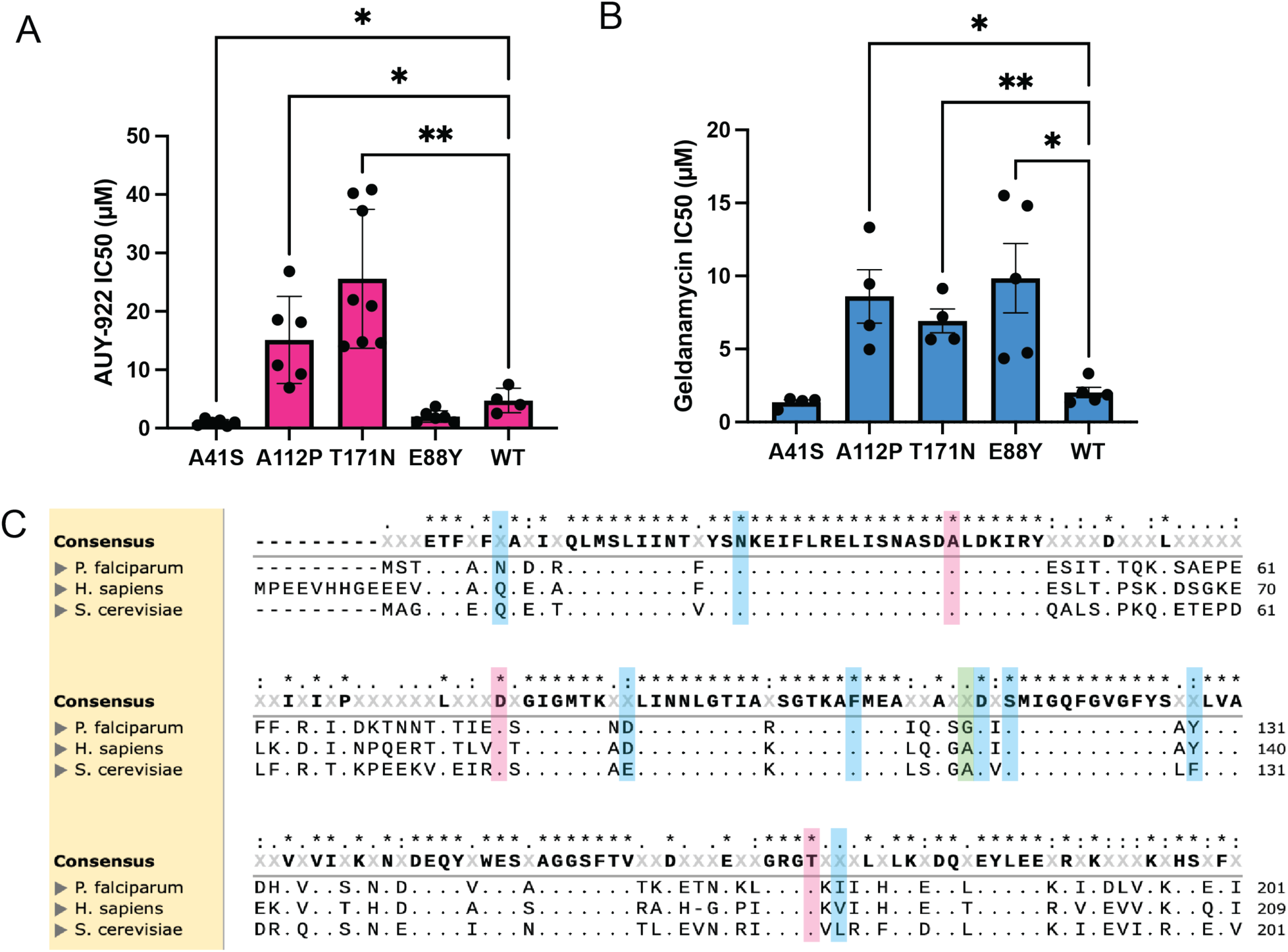
Cross-species comparison of HSP90 inhibitor sensitivity and resistance patterns. **(A)** Cross-resistance analysis of CRISPR-engineered *S. cerevisiae* HSP90 mutations showing IC₅₀ fold-changes against AUY-922 (**B**) Cross-resistance analysis of CRISPR-engineered S. cerevisiae HSP90 mutations showing IC₅₀ fold-changes against geldanamycin. WT represents the wild-type ABC16-Monster strain. **(C)** Multiple sequence alignment of *P. falciparum* HSP90, *S. cerevisiae* HSC82, and human HSP90α. Resistance mutations identified from both *P. falciparum* and *S. cerevisiae* are highlighted: AUY-922 (red), geldanamycin (blue), and shared mutations (green), demonstrating conservation of key residues across species.

The nucleotide binding domain for HSP90 is highly conserved, showing 70% identity between *S. cerevisiae* Hsc82 and *P. falciparum* HSP90, through the first 211 residues, with similar conservation observed with human HSP90α (Figure 3C). Notably, AUY-922 resistance mutations are located at more highly conserved positions compared to geldanamycin resistance mutations.

### Structural Characterization of Resistance Mutations and Drug-Target Interactions

To understand the structural basis of resistance, we performed in silico analysis of the identified mutations using structural models of PfHSP90. The full-length PfHSP90 structure in complex with ADP was generated using AlphaFold3 (Hekkelman et al. 2023), showing the overall protein architecture with the N-terminal ATP-binding domain (Figure 4A, right panel). The left panel shows the crystal structure (3K60) with resistance mutations from both *P. falciparum* and yeast mapped onto the ATP-binding pocket, revealing their spatial distribution relative to the active site (Corbett and Berger 2010). Notably, AUY-922 resistance mutations are located in close proximity to the ATP-binding pocket, while geldanamycin resistance mutations are positioned farther from the active site.

**Figure 4.**
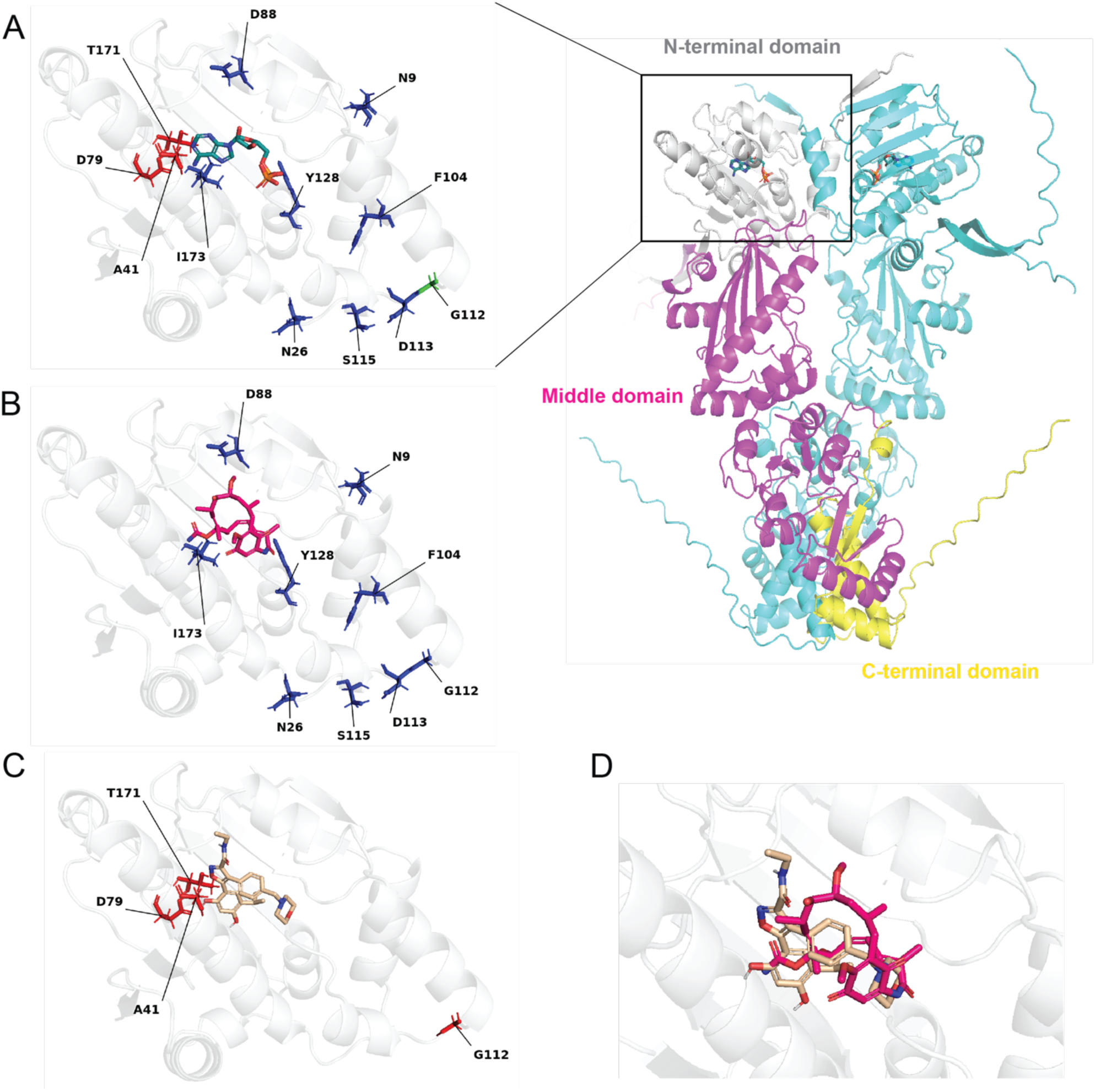
Structural analysis of resistance mutations and ligand interactions. **(A)** Location of resistance mutations in PfHSP90 structure. Left: Crystal structure (3K60) with ADP (turquoise) bound, showing resistance mutations from both *P. falciparum* and *S. cerevisiae* mapped onto the ATP-binding pocket. AUY-922 resistance mutations (red), geldanamycin resistance mutations (blue), and mutations conferring resistance to both inhibitors (green) are indicated. Right: Full-length PfHSP90 homodimer structure in complex with ADP generated using AlphaFold3, showing overall protein architecture with the N-terminal domain (grey), middle domain (magenta), and C-terminal domain (yellow) of one monomer, and the second monomer in cyan. **(B)** Structural context of geldanamycin (magenta) resistance mutations mapped onto the PfHSP90 structure, showing their distribution throughout the N-terminal domain relative to the inhibitor binding site. **(C)** AUY-922 (tan) binding mode in PfHSP90 obtained through computational docking, showing the inhibitor positioned within the ATP-binding pocket. **(D)** Structural overlay demonstrating that both AUY-922 and geldanamycin bind to the same ATP-binding pocket of PfHSP90, with both inhibitors shown in the binding site to illustrate their overlapping binding regions despite different resistance profiles.

For detailed ligand-protein interaction analysis, we employed computational approaches to examine natural substrate and inhibitor binding. ADP binding to PfHSP90 was analyzed using PoseEdit (Diedrich et al. 2023), revealing the extensive interaction network formed by the natural substrate (Figure S4A). ADP forms critical hydrogen bonds with key residues including D79, N37, and F124, establishing the baseline interaction pattern for the ATP-binding pocket. Geldanamycin interactions were modeled through structural alignment with the yeast HSP90-geldanamycin complex, showing that resistance mutations are distributed throughout the N-terminal domain (Figure 4B). PoseEdit analysis revealed the detailed binding interactions of geldanamycin, though this analysis does not include resistance mutations (Figure S4B). Since most geldanamycin resistance mutations are located away from direct contact sites with the inhibitor, they likely affect binding through indirect mechanisms such as altering pocket conformation or dynamics. AUY-922 binding interactions were analyzed through computational docking followed by PoseEdit analysis (Figure 4C, S4C). The analysis revealed that AUY-922 resistance mutations are positioned near the bound inhibitor.

Structural overlay analysis demonstrated that both AUY-922 and geldanamycin bind to the same ATP-binding pocket of PfHSP90, confirming their shared target site despite their different resistance profiles (Figure 4D). This analysis revealed the spatial relationship between the two inhibitors within the binding pocket and provided insights into how their different binding modes may contribute to distinct resistance mechanisms. These structural insights provide a molecular basis for understanding the different resistance profiles observed for these compounds, with AUY-922 resistance mutations directly impacting both inhibitor binding and essential ATP/ADP interactions, explaining the higher barrier to resistance development. These findings align with the substrate envelope hypothesis, which posits that inhibitors mimicking the natural substrate binding mode can minimize the probability of resistance (Özen and Schiffer 2017).

### Binding Affinity Analysis Reveals Enhanced AUY-922 Interactions with A41S Mutant

To validate the binding affinity differences between wild-type and mutant PfHSP90, we expressed and purified the N-terminal domains of PfHSP90 wild-type, PfHSP90 A41S, and PfGRP94 in *E. coli* using Ni-NTA chromatography. Protein purity was confirmed by SDS-PAGE and Western blot analysis (Figure S5), demonstrating successful expression and purification of all three proteins.

We employed fluorescence polarization (FP) to measure competitive binding affinities. This technique exploits the principle that when a fluorescently labeled ligand binds to a target protein, its rotational diffusion becomes restricted, resulting in increased polarization of the emitted light (Figure 5A). We used FITC-labeled geldanamycin (GA-FITC) as the tracer, and the magnitude of polarization change directly reflects the fraction of bound ligand, providing a sensitive assessment of binding affinity. In competitive binding assays, unlabeled inhibitors (geldanamycin or AUY-922) competed with GA-FITC for binding to PfHPS90, causing concentration-dependent reductions in polarization that were fitted to derive Ki values (Kim et al. 2004).

**Figure 5.**
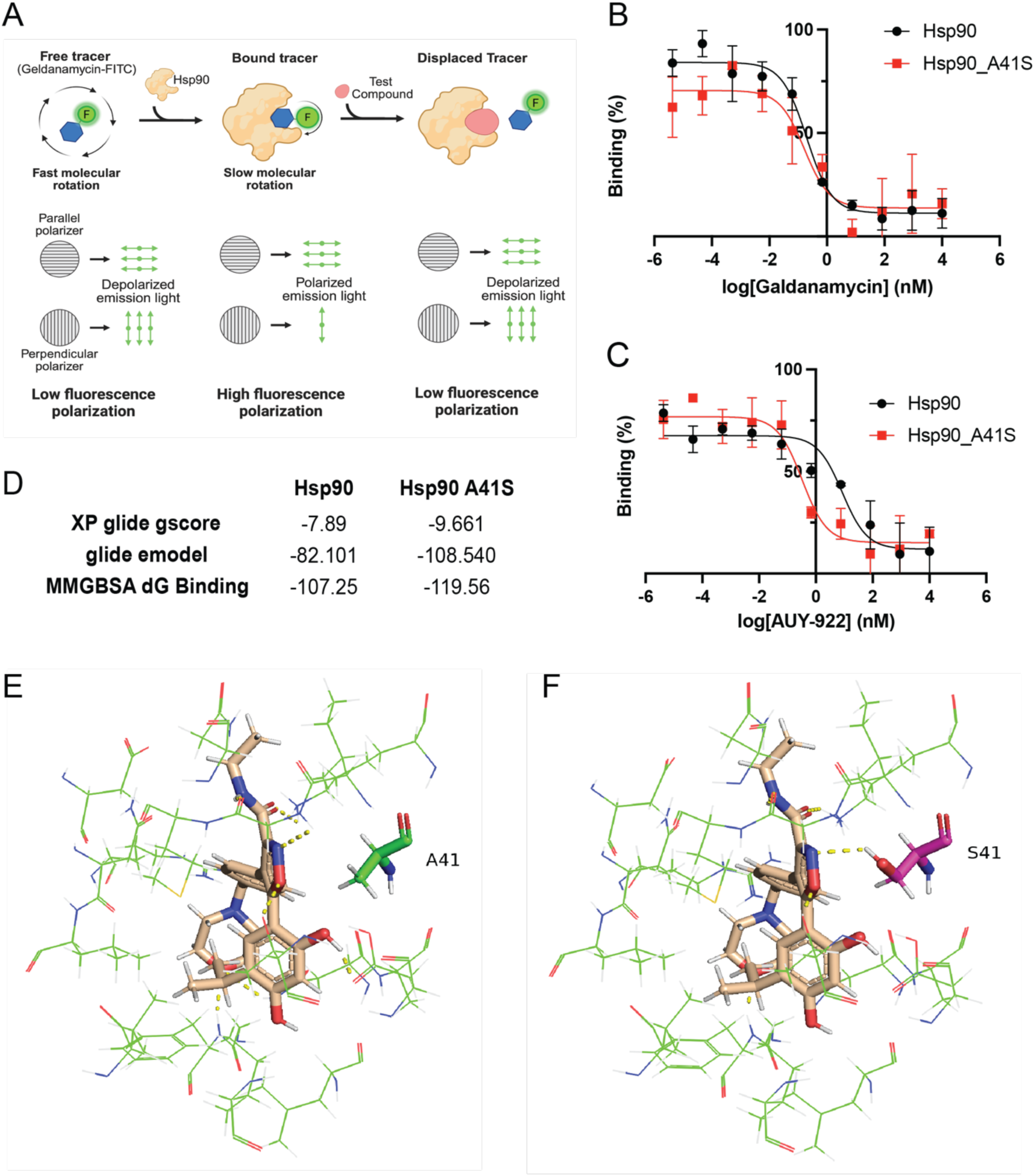
Characterization of binding affinities between PfHSP90 variants and inhibitors. **(A)** Schematic of fluorescence polarization assay principle showing free, bound, and displaced GA-FITC tracer states, illustrating how competitive binding reduces polarization in a concentration-dependent manner. **(B)** Competitive binding curves for geldanamycin showing similar binding profiles between wild-type PfHSP90 (black) and A41S mutant (red), indicating unchanged binding affinity. **(C)** AUY-922 competitive binding curves demonstrating enhanced binding affinity for PfHSP90 A41S mutant (red) compared to wild-type PfHSP90 (black). **(D)** Computational binding analysis showing improved docking scores and binding energies for the A41S mutant across multiple metrics (XP Glide gscore, Glide emodel, and MM-GBSA). **(E)** Structural model of wild-type PfHSP90 with AUY-922 showing the A41 residue (green) position relative to the bound inhibitor. **(F)** Structural model of PfHSP90 A41S mutant with AUY-922 highlighting the S41 residue (magenta) and the additional hydrogen bonding opportunity (yellow dashed line) it provides.

Our results revealed striking differences in binding behavior between the compounds and protein variants. For geldanamycin competition, binding affinity remained essentially unchanged between wild-type PfHSP90 and the A41S variant (Figure 5B), consistent with our cellular IC₅₀ data showing that A41S does not confer resistance to geldanamycin. Note that PfGRP94 showed no detectable binding with GA-FITC under our assay conditions, precluding competitive binding analysis for this protein. In contrast, AUY-922 competition revealed enhanced binding affinity for the A41S variant compared to wild-type protein (Figure 5C). This finding is particularly intriguing because the A41S mutation, which confers resistance to AUY-922 in live parasites, actually increases drug binding affinity—contrary to traditional resistance mutations that typically decrease drug-target interactions. Computational analysis strongly supported this biochemical finding (Figure 5D). All computational metrics—XP Glide gscore (Friesner et al. 2004), Glide emodel, and MM-GBSA binding free energy (Genheden and Ryde 2015)—consistently demonstrated enhanced binding interactions for the A41S mutant compared to wild-type protein, corroborating the experimental fluorescence polarization results. Structural analysis revealed the molecular basis for this enhanced binding: while the wild-type A41 residue shows no direct hydrogen bonding with AUY-922 (Figure 5E, S4C), the A41S mutant provided a strong improvement in docking interactions (Figure 5F, S4D). The S41 residue contributed new hydrogen bonding interactions that were absent in the wild-type protein. This paradoxical enhancement of binding affinity in a resistant mutant represents a fundamentally different resistance mechanism compared to traditional mutations that reduce drug binding affinity.

### HSP90 Knockdown Analysis Reveals Contrasting Drug Mechanisms

To further understand the role of HSP90 as a drug target, we performed conditional knockdowns of HSP90 using the PfDOZI-TetR system previously described (Ganesan et al. 2016; Nasamu et al. 2021). This system enables translational regulation by incorporating 10 TetR-binding aptamer sequences after the HSP90 stop codon. We engineered a vector containing these aptamers, along with an HSP90 epitope tag, a PfDOZI-TetR expression cassette, and a blasticidin-selectable (bsd) marker (Figure 6A). Using CRISPR/Cas9, we integrated this construct into the *P. falciparum* genome. In this system, PfDOZI-TetR binds to the aptamers, resulting in HSP90 translational repression. This repression is relieved by adding anhydrotetracycline (aTc), which prevents PfDOZI-TetR from binding to the aptamers. Given HSP90’s essential role in parasite survival, we carefully calibrated the system to achieve partial protein depletion without catastrophic effects on parasite viability. We determined that an aTc concentration of 5 nM provided an optimal balance (Figure 5B), allowing us to examine drug-target interactions while maintaining sufficient HSP90 function for parasite growth.

**Figure 6.**
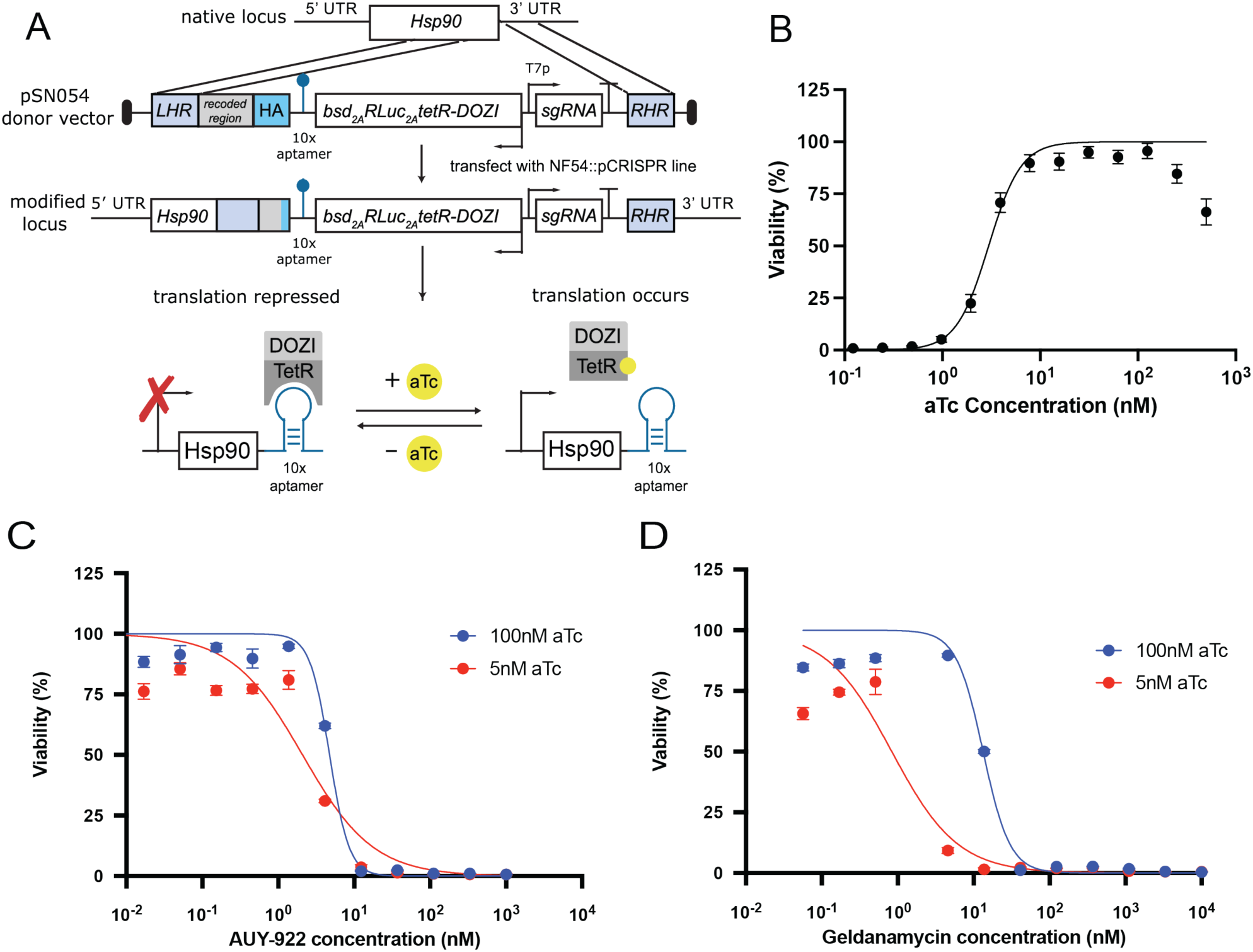
HSP90 conditional knockdown reveals differential dependency of inhibitor activity. **(A)** Schematic representation of the PfDOZI-TetR conditional knockdown system, showing the integration of TetR-binding aptamer sequences, HSP90 epitope tag, PfDOZI-TetR expression cassette, and selection marker. **(B)** Dose-dependent regulation of HSP90 protein levels by varying concentrations of anhydrotetracycline (aTc). Optimization to achieve partial protein depletion while maintaining parasite viability. **(C)** IC_50_ shift assay for AUY-922 under HSP90 knockdown conditions comparing 5 nM vs 100 nM aTc treatment. **(D)** IC_50_ shift assay for geldanamycin under HSP90 knockdown conditions comparing 5 nM vs 100 nM aTc treatment.

Under these conditions, we tested our conditional parasites against HSP90 inhibitors and control compounds. The results revealed striking differences between the compounds. AUY-922 showed almost no IC₅₀ shift under HSP90 depletion conditions (Figure 6C), with similar IC₅₀ values at both 100 nM and 5 nM aTc concentrations. In contrast, the conditional knockdown rendered parasites dramatically more sensitive to geldanamycin (Figure 6D), with a substantial leftward shift in the dose-response curve under HSP90 depletion (5 nM aTc) compared to the control condition (100 nM aTc). Similar patterns were observed with radicicol, another HSP90 inhibitor, which showed increased sensitivity under HSP90 knockdown conditions (Figure S6A). As a specificity control, we tested GNF179, which acts through an unrelated mechanism, and observed minimal difference between the two aTc conditions (Figure S6B), confirming that the differential effects were specific to HSP90 inhibitors. These data strongly suggest that AUY-922 may exert its antimalarial activity through a different mechanism than direct HSP90 inhibition, while confirming HSP90 as the primary target for geldanamycin and radicicol.

## Discussion

Our findings reveal that AUY-922 exhibits an exceptionally high barrier to resistance development, representing a rare example of a highly resilient antimalarial compound. The stark contrast with geldanamycin—despite both targeting the same HSP90 binding site—challenges assumptions that compounds sharing identical targets exhibit similar resistance profiles. The most striking discovery is the paradoxical enhancement of AUY-922 binding by the single, recovered A41S mutation We hypothesize that this mutation may create a molecular trap that sequesters drug away from critical cellular targets, rather than following the classical competitive inhibition pattern that typically weakens drug-target interactions. This enhanced binding mechanism may represent an underexplored paradigm in antimalarial drug development. We propose that A41S increases AUY-922 affinity for cytosolic PfHSP90 while reducing availability for other essential HSP90 homologs. This redistribution protects the critical family members required for antiparasitic activity, conferring resistance despite stronger binding at the primary target. Mechanistically analogous phenomena have been observed in other systems where protein-mediated drug sequestration reduces effective target engagement and produces resistance — for example, in cancer cells, LAPTM4B overexpression sequesters doxorubicin and delays nuclear delivery (Li et al. 2010), providing a precedent for protein-facilitated drug trapping as a resistance mechanism.

The differential responses observed in our conditional knockdown experiments—where HSP90 depletion minimally affected AUY-922 sensitivity but significantly increased sensitivity to geldanamycin and radicicol—point toward distinct mechanisms of action for these structurally different HSP90 inhibitors. We propose a multi-target model where the antimalarial activity of AUY-922 depends on coordinated inhibition of multiple HSP90 family members, while geldanamycin acts primarily through cytosolic HSP90. This hypothesis is supported by previous studies showing AUY-922’s broader binding profile across HSP90 isoforms compared to geldanamycin’s more selective targeting (Murillo-Solano et al. 2017). The cross-species differences in A41S mutation effects further support this multi-target hypothesis. While A41S confers resistance in *P. falciparum* but increases sensitivity in *S. cerevisiae*, this opposing effect correlates with fundamental differences in HSP90 family complexity between the two organisms. *P. falciparum* expresses four HSP90 homologs across different cellular compartments—PfHSP90 (cytosolic), PfGRP94 (ER), PfTRAP1 (mitochondrial), and PfHSP90-A (apicoplast)—while *S. cerevisiae* has only two cytosolic variants: constitutively expressed HSC82 and stress-induced HSP82. The A41S mutation may redistribute AUY-922 binding in favor of cytosolic PfHSP90, thereby protecting other essential family members from inhibition in the more complex *P. falciparum* system. Among these potential protected targets, PfGrp94 has emerged as a promising antimalarial drug target, with recent studies demonstrating its druggability and essential roles in parasite protein export and survival (Muzenda et al. 2025; Stofberg et al. 2024).

Emerging evidence highlights the attractiveness of developing “irresistible” antimalarial compounds that exhibit exceptionally high barriers to resistance (Yang et al. 2022). One potential mechanism for achieving such resistance barriers is through multi-target engagement. A compelling example of this principle is WM382, a dual plasmepsin IX/X antimalarial inhibitor that demonstrates an MIR value >10⁹ compared to single-target plasmepsin inhibitors, presumably because resistance would require simultaneous modifications to both target enzymes (De Lera Ruiz et al. 2022). Similarly, the vinyl sulfone WLL exhibits exceptionally low resistance propensity through dual targeting of proteasome β2 and β5 subunits (Deni et al. 2023). The exceptionally high MIR value observed for AUY-922 aligns with this principle, suggesting that effective antimalarial activity requires coordinated inhibition of multiple HSP90 family members.

Traditional drug discovery focuses on optimizing compounds against individual targets, but our findings suggest that deliberately designing compounds to engage multiple essential proteins within a family may provide superior resistance barriers. This principle aligns with broader polypharmacological strategies, where multi-target drugs have demonstrated superior resistance barriers across diverse therapeutic areas (Makhoba et al. 2020). This multi-target strategy extends beyond protein families to encompass different proteins within the same pathway, as exemplified by the antimalarial RYL-581, which simultaneously targets three distinct binding sites across separate mitochondrial respiratory chain proteins (Yang et al. 2021). However, this approach must be balanced against potential safety concerns. For infectious disease drugs, selectivity for pathogen targets over host proteins is crucial to minimize toxicity (Lawong et al. 2024; Mansfield et al. 2024; Tye et al. 2022), and multi-target engagement could potentially increase the risk of host cell toxicity through broader effects on essential cellular proteins.

Our findings have important implications for antimalarial drug development. The contrasting resistance profiles of two compounds targeting the same binding site demonstrate that resistance risk cannot be predicted solely from target identity or binding mode. The identification of AUY-922’s resistance profile provides a framework for recognizing and developing compounds with enhanced durability against resistance. Understanding how to optimize compounds that resist evolutionary escape represents a fundamental advance in medicinal chemistry that could significantly extend the useful lifespan of therapeutic interventions, contributing to more sustainable malaria control strategies and broader efforts to combat drug resistance across diverse disease contexts.

## Material and Method

### P. falciparum culturing

*P. falciparum* Dd2 strain parasites were cultured under standard conditions^5^, using RPMI media supplemented with 0.05 mg/ml gentamycin, 0.014 mg/ml hypoxanthine (prepared fresh), 38.4 mM HEPES, 0.2% sodium bicarbonate, 3.4 mM sodium hydroxide, 0.05% O^+^ Human serum (denatured at 56^0^C for 40 min -Interstate Blood Bank, Memphis, TN), and 0.0025% albumax. Human O^+^ whole blood was obtained from TSRI Normal Blood Donor Services (La Jolla, CA). Leukocyte-free erythrocytes were stored at 50% hematocrit in RPMI-1640 screening media (as above, but without O^+^ human serum and with double albumax concentration) at 4^0^C for one to three weeks before experimental use. Cultures were monitored every one to two days via direct observation of parasite infection using light microscopy-based observation of Giemsa-stained thin blood smears of parasite cultures.

### Drug preparation

The drug stock solutions were prepared by dissolving the compounds in dimethyl sulfoxide (DMSO) to achieve concentrations of 100 mM. Aliquots of 10 mM working stocks were prepared from these stock solutions and stored at −20°C for long-term storage. In all experiments, the final concentration of DMSO was maintained below 0.5% to minimize potential solvent effects.

### *In vitro* drug sensitivity and IC_50_ determinations

*In vitro* drug susceptibility was determined by quantifying the total parasitic DNA content using SYBR Green staining in a 384-well plate format. Synchronized ring-stage *P. falciparum* Dd2-B2 clone parasites were cultured in the presence of 12-point, 2-fold serial dilutions of the test compounds in black, clear-bottom plates. An artemisinin dilution series was conducted in parallel as a positive control. After a 72-hour incubation under standard culture conditions, the SYBR Green I fluorescent nucleic acid dye was added to the cultures. The fluorescence signal, which is proportional to the amount of parasitic DNA present, was measured using a Spectramax M5 plate reader (Molecular Devices) with excitation and emission wavelengths of 480 nm and 530 nm, respectively. Following positive control fluorescence subtraction and signal normalization, the IC50 values (effective concentration for 50% growth inhibition) were calculated using the Levenberg-Marquardt algorithm implemented in the Collaborative Drug Discovery database (Burlingame, CA. www.collaborativedrug.com).

### Liver stage assay

Compounds were dissolved in DMSO (10 mM), and 10 nL of each compound (0.1% final DMSO concentration per well) was transferred using an ECHO 650 (Beckman) into the assay plates within the range of 10 µM to 0.5 nM atovaquone (1 µM) and 0.1% DMSO served as the positive and negative controls, respectively. Human hepatic cells (3 × 10³; HepG2-A16-CD81-EGFP) in 5 µL of DMEM medium (2 × 10⁵ cells/mL, 5% FBS, 5X Pen/Strep/Glu) were seeded in 1536-well plates (Greiner BioOne) 20 hours prior to the actual infection. PbLuc sporozoites were dissected from the salivary glands of infected Anopheles stephensi mosquitoes and filtered twice through a 20 µm nylon pore cell strainer (Swann et al. 2016). *Plasmodium berghei* luciferase (PbLuc) sporozoites were obtained by dissecting salivary glands of infected Anopheles stephensi mosquitoes provided by The SporoCore, University of Georgia, GA, USA (SporoCore.uga.edu). The sporozoites were resuspended in screening media, counted using a hemocytometer, and their final concentration was adjusted to 200 sporozoites per µL. The HepG2 cells were then infected with 1,000 sporozoites per well (5 µL), and the plates were spun down at 37°C for 3 minutes in an Eppendorf 5810 R centrifuge with a centrifugal force of 330 RCF on the lowest acceleration and brake settings. After incubation at 37°C in 5% CO₂ for 48 hours, the media was removed by spinning the inverted plates at 130 RCF for 1 minute. 2 µL of Bright-Glo™ Luciferase Assay System (Promega) was dispensed into the wells. Immediately after the addition of the Bright-Glo reagent, the plates were read by the Pherastar FSX reader (BMG Labtech). Luminescence intensity values were normalized against the positive (Atovaquone) and negative (DMSO) controls. All experiments were conducted in four technical replicates and repeated three times.

### Cytotoxicity assay

Plates were prepared according to the liver stage conditions, except 5 µl of screening media was used instead of sporozoites (PMID: 27275010). After incubation at 37°C for 48 hours, the media was removed by spinning the inverted plates at 130 RCF for 1 minute. Then, 2 µl of CellTiter-Glo® Luminescent Cell Viability Assay (Promega) was dispensed into each well. Immediately after adding the CellTiter-Glo reagent, the plates were read using the Pherastar FSX reader (BMG Labtech). Luminescence intensity values were normalized against positive (Puromycin) and negative (DMSO) controls. IC_50_ values were calculated in CDD Vault (Burlingame, CA). All experiments were conducted in four technical replicates and repeated at least two times.

### Determination of parasite stage by flow cytometry

*P. falciparum* cultures were first synchronized using 5% w/v sorbitol solution to obtain a population of ring-stage parasites. The synchronized cultures were then adjusted to 2% hematocrit and 2% parasitemia in T12.5 flasks. These cultures were treated with either 5 × IC_50_ of AUY-922, 5 × IC_50_ of geldanamycin, or an equivalent volume of DMSO as a control. The treated cultures were incubated for a total of 48 hours under standard culture conditions. At 24- and 48-hours post-treatment, samples were collected for flow cytometric analysis. The parasites were stained with 2μM SYTO61, a red fluorescent nucleic acid stain, for 20 minutes at room temperature. Following staining, the samples were washed with PBS to remove excess dye. Flow cytometry was then performed to analyze the DNA content and subsequently determine the distribution of parasite stages in each treatment condition.

### In vitro resistance evolution in *Plasmodium*

Independent *P. falciparum* Dd2 clones were subjected to gradually increasing concentrations of AUY-922 over 44 weeks or geldanamycin over 8 weeks, starting at IC30 concentrations. The parasitemia levels were monitored daily by visual inspection using light microscopy of Giemsa-stained thin blood smears prepared from the parasite-infected erythrocyte cultures. Resistance was defined as achieving an IC50 value (the concentration for 50% growth inhibition) at least 2-fold higher than that of the untreated parental Dd2 strain. We also examined geldanamycin resistance using an early cloning protocol as previously described (Lim et al. 2016), where parasites were treated with 3× IC90 concentrations for 96 hours, then seeded in 96-well plates and monitored for growth recovery. Upon attaining resistance phenotypes, genomic DNA was extracted from both the resistant cultures and the parental line for whole-genome sequencing analysis or PCR amplification of the PfHSP90 DNA sequence, which was then sent for Nanopore sequencing (Plasmidsaurus).

### Minimum inoculum of resistance (MIR)

The MIR assay was performed using single-step continuous pressure of 3×IC₉₀ drug concentrations for geldanamycin and 3.5×IC₅₀ for AUY-922 and the DSM265 control on two parasite inoculum groups per compound. For AUY-922 and DSM265 control, inoculum sizes were 2.5 × 10⁶ parasites (four wells; 2% hematocrit) and 3 × 10⁷ parasites (three wells; 3% hematocrit). Geldanamycin testing used 3.5 × 10⁶ parasites per well in 24-well plates and 4.8 × 10⁸ parasites per w ell in 6-well plates. Wells were monitored daily by thin blood smear during the initial drug killing phase until parasite clearance. Geldanamycin cultures received additional monitoring by flow cytometry (24-well plates) and Giemsa-stained smears (6-well plates). Following clearance, drug media was changed three times weekly, cultures were passaged weekly by replacing one-third of the culture volume with fresh RBC media and monitored twice weekly by flow cytometry for recrudescence over 60 days. Genomic DNA from resistant cultures and parental lines was extracted for whole-genome sequencing analysis.

### Whole-genome sequencing and analysis

Cultures were initially lysed with 0.15% saponin and washed twice with 1x PBS. Genomic DNA was extracted using the QIAamp DNA Blood Mini Kit (Qiagen) according to the manufacturer’s instructions. Whole-genome sequencing (WGS) libraries were prepared using the Nextera Flex DNA Library Preparation Kit (Illumina) following the manufacturer’s protocol. The libraries were multiplexed and sequenced on an Illumina MiSeq platform to generate 300 bp paired-end reads. The generated sequencing reads were quality-checked using FastQC (version 0.11.9) and trimmed using Trimmomatic (version 0.39) to remove adapters and low-quality bases. The trimmed reads were then aligned to the *P. falciparum* 3D7 reference genome (PlasmoDB, version 48) using the Burrows-Wheeler Aligner (BWA-MEM, version 0.7.17). PCR duplicates were removed using Picard (version 2.23.8), and unmapped reads were filtered out using SAMtools (version 1.10). Base quality score recalibration (BQSR) was performed using the Genome Analysis Toolkit (GATK, version 4.1.8) to adjust for systematic errors in base quality scores. The GATK HaplotypeCaller was then used to identify potential single nucleotide variants (SNVs) in the test parasite lines. The identified SNVs were filtered based on the following quality scores: variant quality as a function of depth (QD) > 1.5, mapping quality (MQ) > 40, minimum base quality score (MBQS) > 18, and read depth (DP) > 5. The resulting high-quality SNVs were annotated using SnpEff (version 4.3t) to predict their functional impact on genes and proteins.

### CRISPR/Cas9 gene editing in *P. falciparum*

To validate the role of the A41S mutation in PfHSP90 in conferring resistance to AUY-922, we performed CRISPR/Cas9-mediated gene editing in the *P. falciparum* Dd2 strain. As a control, we also generated a synonymous mutation at the same position (A41A). The CRISPR/Cas9 plasmid, pDC2-coCas9-gRNA, containing a codon-optimized SpCas9 and a guide RNA (gRNA) cassette, was used for gene editing (Adjalley and Lee 2022). The gRNA targeting the PfHSP90 locus was designed using Benchling (www.benchling.com) and cloned into the *Bbs*I site of the plasmid. The donor repair template, encoding the A41S mutation and silent shield mutations at the gRNA-binding site to prevent re-cutting, was synthesized as a double-stranded DNA fragment (GeneArt, Thermo Fisher Scientific) and cloned into the AatII/EcoRI sites of the plasmid using NEBuilder HiFi DNA Assembly (New England Biolabs). Successful insertion of the gRNA and donor template was confirmed by Sanger sequencing.

Synchronized ring-stage *P. falciparum* Dd2 parasites at 10% parasitemia were electroporated with 50 µg of the CRISPR/Cas9 plasmid. Parasites were resuspended in cytomix (120 mM KCl, 0.2 mM CaCl_2_, 2 mM EGTA, 10 mM MgCl_2_, 25 mM HEPES, 5 mM K2HPO4, 5 mM KH2PO4; pH 7.6) to a final volume of 420 µL and electroporated using a Bio-Rad Gene Pulser II (0.31 kV, 950 µF). After a 1-hour recovery, the electroporated parasites were washed and cultured in complete medium at 3% hematocrit. Drug selection with 5 nM WR99210 (Jacobus Pharmaceuticals) was applied 24 hours post-electroporation and maintained until day 8. Surviving parasites were then cultured without drug pressure, and clonal populations were obtained by limiting dilution. Genomic DNA was extracted from the clonal parasite lines using the QIAamp DNA Blood Mini Kit (Qiagen), and the PfHSP90 locus was PCR-amplified using primers flanking the edited region. The presence of the desired mutations was confirmed by Sanger sequencing.

### Antimalarial Resistome Barcode (AReBar) assay for target deconvolution

To investigate the impact of HSP90 inhibitors on different *P. falciparum* drug resistant lines (Supplemental Table 3), we employed a competitive growth assay using 52 DNA barcoded lines covering >30 modes of action, essentially as previously described (Carrasquilla et al. 2022). The pooled culture was divided into three conditions: untreated control, AUY-922 at 3 × IC_50_, and geldanamycin at 3 × IC_50_. Cultures were maintained at 2% hematocrit and 1-5% parasitemia for 14-18 days, with media and compound replaced every 48 hours. Samples were collected at day 0 and day 14, and the barcode region was amplified from saponin-lysed pellets using primers p1356 and p1357 (Supplemental Table 1) for sequencing using the minION (Mk1d) platform. Sequencing data was processed to determine the relative proportion of each barcode over time, normalized to its initial proportion, and log2-transformed. Differentially represented barcodes under HSP90 inhibitor treatment compared to the untreated control were identified using the DESeq2 R package (Love et al. 2014).

### Yeast strain and culture conditions

The drug-sensitive ABC16-Green Monster strain, which lacks 16 ABC transporter genes, was used for all yeast experiments (Suzuki et al. 2011). Yeast cells were cultured in YPD media overnight at 30°C with shaking at 250 RPM.

### CRISPR-Cas9 allelic replacements in *S. cerevisiae*

CRISPR/Cas9 vectors p414 and p426 from the Church lab (Addgene) (DiCarlo et al. 2013) were modified for compatibility with the ABC16-Monster strain. The Trp selection marker of p414 was replaced with met15, and the Ura selection marker of p426 was replaced with leu2. Cas9-expressing ABC16-Monster yeast cells were generated by transforming the met15-modified p414 plasmid using the standard lithium acetate method and selecting on CM-glucose plates lacking methionine. All DNA sequences were amplified using Q5 High-Fidelity DNA Polymerase (New England Biolabs). Donor DNA containing the desired allelic replacements was synthesized as gBlocks Gene Fragments (Integrated DNA Technologies). The p426 plasmid with the desired gRNA sequence was constructed using a three-fragment HiFi assembly. PCR products were treated with DpnI (New England Biolabs) at 37°C for 1 hour, followed by heat inactivation at 80°C for 20 minutes. The assembly reaction was performed at 50°C for 30 minutes, and the resulting product was transformed into 5-alpha Competent E. coli (New England Biolabs). Cas9-expressing ABC16-Monster yeast cells were transformed with the gRNA plasmid and donor DNA fragments, and transformants were selected on CSM plates lacking methionine and leucine. Genomic DNA from single colonies was isolated, and the presence of the desired mutations was initially confirmed by Sanger sequencing (Eton Bioscience).

### *S. cerevisiae* dose-response assays

To evaluate compound activity against yeast cells, single colonies of the ABC16-Monster strain harboring the desired mutations were inoculated into 3 mL of YPD media and cultured overnight at 30°C with shaking. The following day, cultures were diluted to log phase, and 50 µL of the diluted cultures were dispensed into the wells of a 384-well plate. Nine 1:2 serial dilutions of each compound were prepared in biological duplicates, resulting in final concentrations ranging from 0.005 to 100 µM. The plate was incubated at 30°C for 18 hours, and the optical density at 600 nm (OD_600_) was measured. IC_50_ values were calculated by subtracting the OD_600_ values of positive controls (cells treated with 250 µM fluconazole) and performing nonlinear regression on [inhibitor] vs. response with a variable slope using GraphPad Prism. As a negative control, parallel cultures without compound treatment were included.

### Computational modeling and docking analysis

The Schrodinger Maestro small molecule modeling suite^XX^ was used for all modeling performed. The PfHSP90 crystal structure (PDB ID 3K60, PF3D7_0708400, cytosolic PfHSP90) from the Protein Data Bank was used for docking AUY-922 and modeling the *P. falciparum* A41S mutant. The Schrödinger Prime package was used to create the mutant residue and select the optimal sidechain conformation within the binding pocket. Prime minimization was performed to determine a low-energy protein conformation with minimal deviation from the starting X-ray coordinates. The N-terminal domain containing the ATP-binding pocket was isolated from each structure for docking analysis. Protein structures were prepared using the Protein Preparation Wizard and refined to alleviate steric clashes and structural strain. The AUY-922 ligand was prepared using LigPrep with default parameters. Glide XP (extra precision) docking was employed to dock AUY-922 into all three binding sites. The resulting docked poses were analyzed using the Prime MM-GBSA workflow to calculate binding free energies (ΔG) and estimate relative binding affinities. For geldanamycin analysis, we aligned the 3K60 structure with the yeast HSP90 structure in complex with geldanamycin (PDB: 1A4H). The full-length PfHSP90 structure in complex with ADP was generated using AlphaFold3. Final docked poses were exported to PyMOL for structural visualization, analysis, and figure preparation.

### Protein construct and purification

The N-terminal domains of *P. falciparum* HSP90 and GRP94, each containing a C-terminal hexa-histidine tag, were cloned into the pET21 vector and transformed into NiCo21(DE3) competent E. coli cells. Successful transformants were grown overnight at 37°C in LB medium supplemented with 100 µg/mL ampicillin. For protein expression, 200 mL LB cultures containing ampicillin were inoculated and grown at 37°C until OD reached 0.6. Protein expression was induced with 1 mM IPTG and cultures were incubated overnight at 16°C. Cells were harvested by centrifugation at 4,000 × g for 20 minutes at 4°C and resuspended in lysis buffer (5% glycerol, 0.3 M NaCl, 20 mM HEPES, 30 mM imidazole-HCl, 0.5% CHAPS, 21 mM MgCl₂, 2% CHAPS, 0.1 mg/mL lysozyme, 0.5 U/mL benzonase, and Protease Inhibitor Cocktail). After sonication and clarification at 12,000 × g for 20 minutes, the supernatant was purified using Ni-NTA chromatography, washed extensively (5% glycerol, 0.3 M NaCl, 20 mM HEPES, 30 mM imidazole-HCl, and 21 mM MgCl₂), and eluted with a stepwise imidazole gradient (150-350 mM). Purified proteins were concentrated using 10 kDa MWCO Amicon filters and buffer exchanged into 50 mM HEPES (pH 7.5), 500 mM NaCl. Protein purity was assessed by SDS-PAGE and concentrations were determined using the Qubit Protein Assay Kit.

### Fluorescence polarization binding assay

Binding affinities between HSP90 inhibitors and purified PfHSP90 or PfHSP90-A41S were determined using FITC-labeled geldanamycin (GA-FITC) in a competition assay format. Initial direct binding experiments with serial dilutions of the purified proteins and GA-FITC in HSP90 assay buffer (BPS Bioscience) containing 2 mg/ml BSA and 2 mM DTT established optimal conditions. The Kd values from these experiments guided the selection of protein (10 nM) and GA-FITC (1 nM) concentrations for competition assays. Test compounds were serially diluted in DMSO starting from 1 mM with eight 10 × dilutions and added to the protein/GA-FITC mixture. Assays were performed in 384-well black, low-binding microplates with 50 μl final volume. After 16 hours incubation at 4 °C with gentle shaking, fluorescence polarization was measured using a PHERAstar FSX plate reader (excitation 485 nm, emission 530 nm).

### Generation of cKD parasite lines

To investigate the role of HSP90 as a drug target, we used CRISPR-Cas9 to generate parasite cell lines stably expressing the TetR-DOZI-RNA aptamer module for conditional regulation of HSP90 expression. These transgenic lines also contained the reporter construct *Renilla* luciferase (RLuc) and the selection marker BSD. To construct the donor plasmids, PCR-amplified right homology regions and synthesized DNA fragments corresponding to the left homology regions fused to the recodonized 3′-end of the HSP90 gene with 10× TetR-binding aptamers inserted at the 3′-end, as well as the target-specifying gRNA sequences, were cloned via Gibson assembly into the pSN054 linear vector. The donor vector enables fusion of 2x hemagglutinin tag to the protein, but this approach did not yield viable parasites, therefore the final edited cell line was not tagged. The final constructs were confirmed by restriction digests and Sanger sequencing.

Transfections into Cas9- and T7 RNA polymerase–expressing NF54 parasites were carried out by preloading erythrocytes with the donor plasmids as described previously. Cultures were maintained in 500 nM anhydrotetracycline (aTc; Sigma-Aldrich, 37919) and blasticidin-S (2.5 μg/ml; RPI Corp B12150-0.1). Parasite cell lines stably integrating the donor plasmids were monitored via Giemsa smears and RLuc measurements.

### cKD viability assays

To assess the effect of conditionally perturbing HSP90 expression on drug sensitivity, parasites were washed three times with media and then cultured in the presence of aTc (100 nM and 5 nM) at 2% hematocrit and 0.5% parasitemia for 72 hours. Drug sensitivity assays were performed in triplicate using 384-well plates, and luminescence signals were measured using either PHERAstar FSX or a GloMax (Promega) plate reader following the Renilla-Glo® Luciferase Assay System (Promega) manual.

### Statistical analysis

All statistical analyses were performed using GraphPad Prism software. For comparisons involving three or more groups, two-way ANOVA was performed followed by unpaired t-test with Welch’s correction. When comparing two groups, an unpaired t-test with Welch’s correction was used. Statistical significance was defined as follows: * P ≤ 0.05, ** P ≤ 0.01, *** P ≤ 0.001 and **** P ≤ 0.0001. All data are presented as mean ± standard error of the mean (SEM). All experiments were repeated at least three times.

## Acknowledgements

Partial support for this work was provided by the Gates Foundation (INV-039628 to EAW; INV-033538 to DAF and EAW). EAW acknowledges support from the NIH (R01 AI169892). DAF also acknowledges support from the NIH (R01 AI185559).

## Author contributions

Conceptualization: EAW, F-HK. Methodology: F-HK, EAW. Investigation: F-HK, AKL, MLS, AF, JH, TN, JO, NB, KK, GG, CP, LG, SO. Formal analysis: F-HK. Whole genome sequencing analysis: TY, HP, A-CU. In silico analysis: GLD. Data curation: F-HK. Resources: EAW, DAF, AC. Supervision: EAW, JN, ML, DAF, DFW. Writing – original draft: F-HK. Writing – review & editing: F-HK, EAW, TN, JO, NB, KK, ML, CP, LG, JN, SO, GLD, DFW.

---

**Figure S1.**
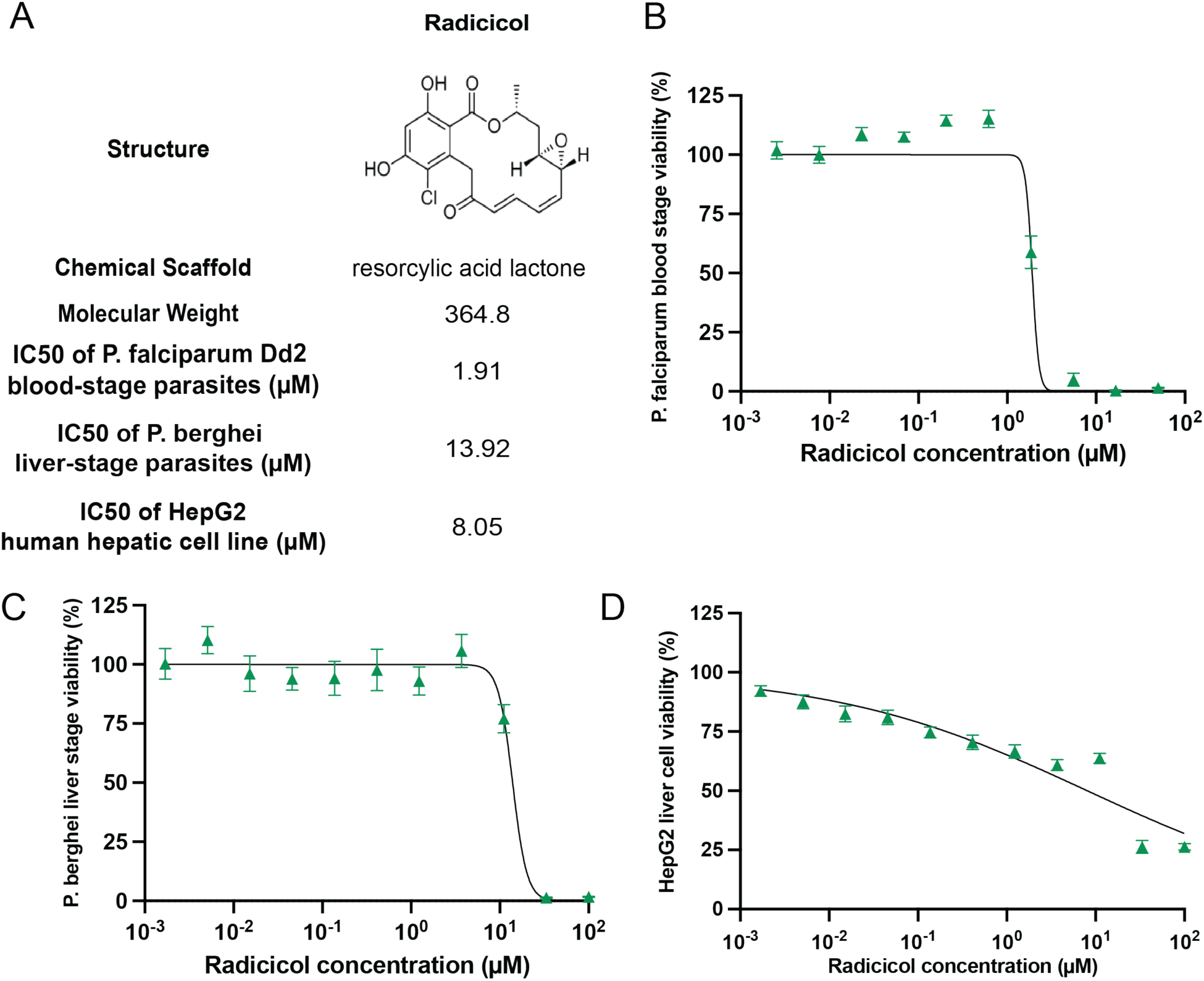
Antimalarial characterization of radicicol as an HSP90 inhibitor. **(A)** Chemical structure and biological activity profile of radicicol showing its resorcylic acid lactone scaffold and IC₅₀ values against different parasite stages and human cells. **(B)** Dose-response curve of radicicol against *P. falciparum* Dd2 blood-stage parasites. **(C)** Dose-response curve of radicicol against *P. berghei* liver-stage parasites. **(D)** Cytotoxicity assessment of radicicol in HepG2 human hepatocytes.

**Figure S2.**
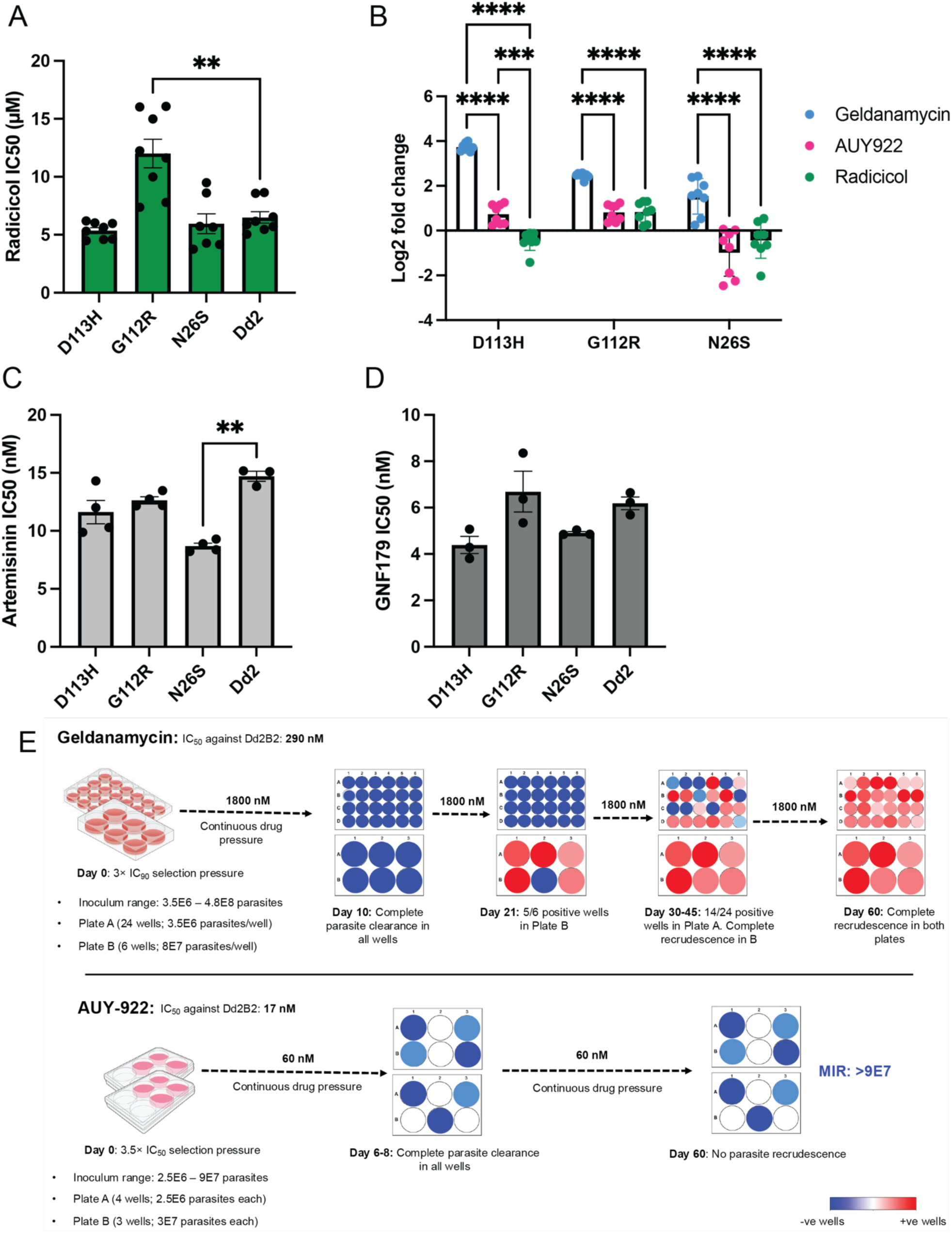
Cross-resistance characterization of geldanamycin-resistant clones and Minimum Inoculum for Resistance analysis. **(A)** IC₅₀ values of geldanamycin-resistant clones against radicicol. (B) Log₂ fold-change analysis of geldanamycin-resistant mutations (D113H, G112R, N26S) against geldanamycin (blue), AUY-922 (magenta), and radicicol (green). **(C)**IC₅₀ values of geldanamycin-resistant clones against artemisinin. **(D)** IC₅₀ values of geldanamycin-resistant clones against GNF179. **(E)** Minimum Inoculum for Resistance (MIR) assay results. Top: Geldanamycin MIR assay progression from Day 0 to Day 60. Bottom: AUY-922 MIR assay showing no recrudescence (MIR >9×10⁷). Blue circles represent negative wells, red circles represent positive wells, white circles represent wells not tested.

**Figure S3.**
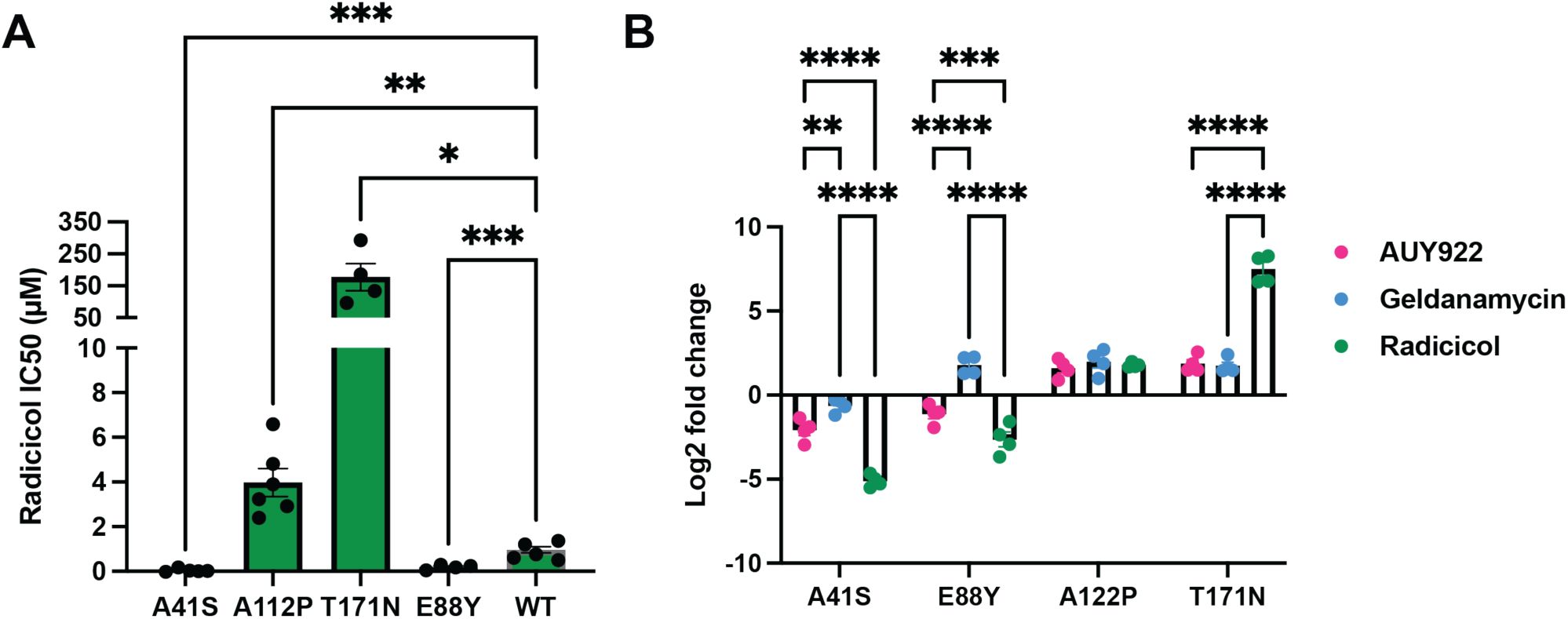
Cross-species comparison of HSP90 inhibitor sensitivity in *S. cerevisiae*. **(A)** IC₅₀ values of CRISPR-engineered *S. cerevisiae* HSP90 mutations against radicicol. **(B)** IC₅₀ fold-changes of CRISPR-engineered *S. cerevisiae* HSP90 mutations against AUY-922, geldanamycin, and radicicol, demonstrating mutation-specific resistance and sensitivity patterns across the three HSP90 inhibitors.

**Figure S4.**
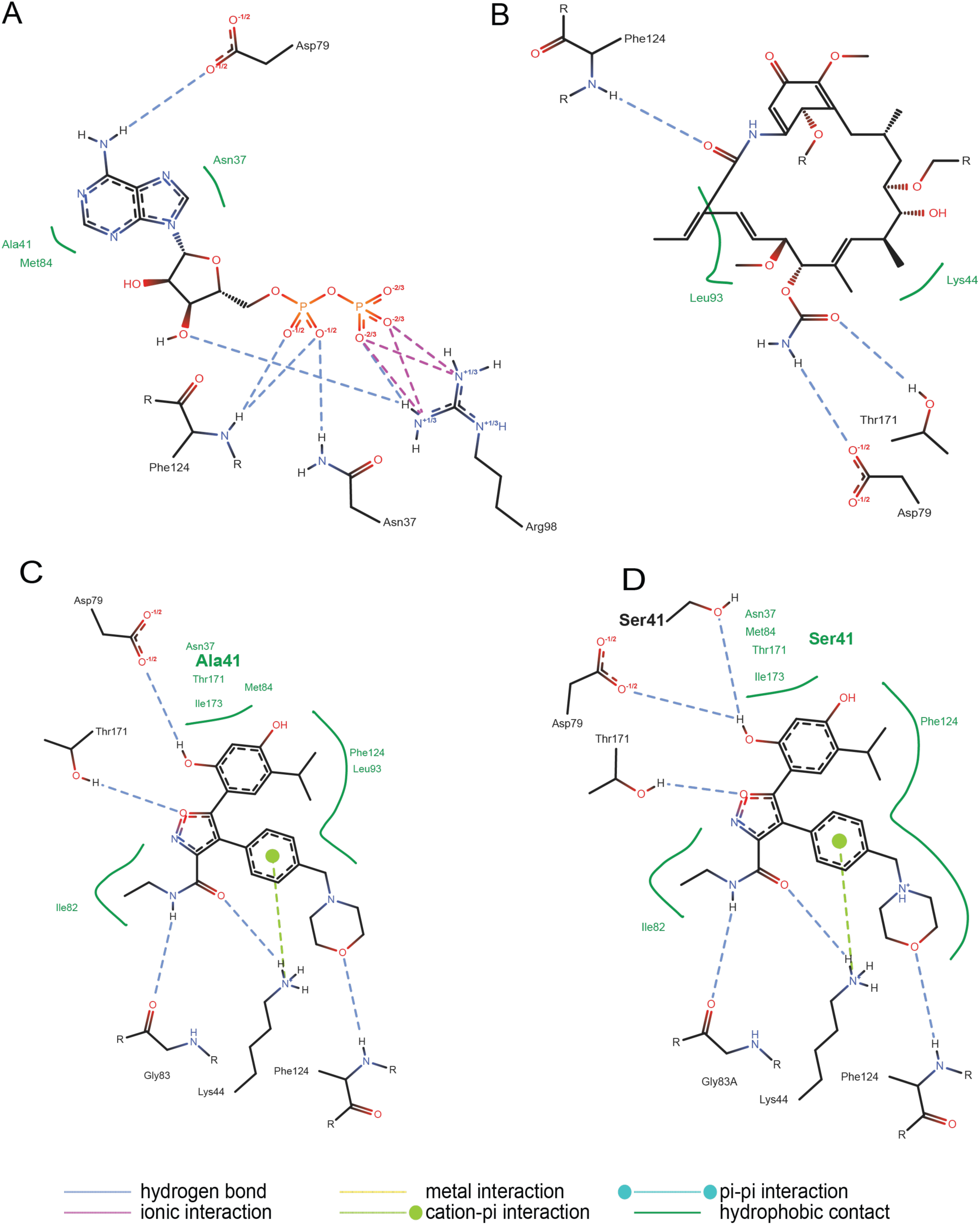
PoseEdit ligand interaction analysis. **(A)** PoseEdit interaction diagram of ADP binding to PfHSP90. **(B)** PoseEdit interaction diagram of geldanamycin binding to PfHSP90. **(C)** PoseEdit interaction diagram of AUY-922 binding to wild-type PfHSP90 with Ala41 highlighted. **(D)**PoseEdit interaction diagram of AUY-922 binding to PfHSP90 A41S mutant with Ser41 highlighted.

**Figure S5.**
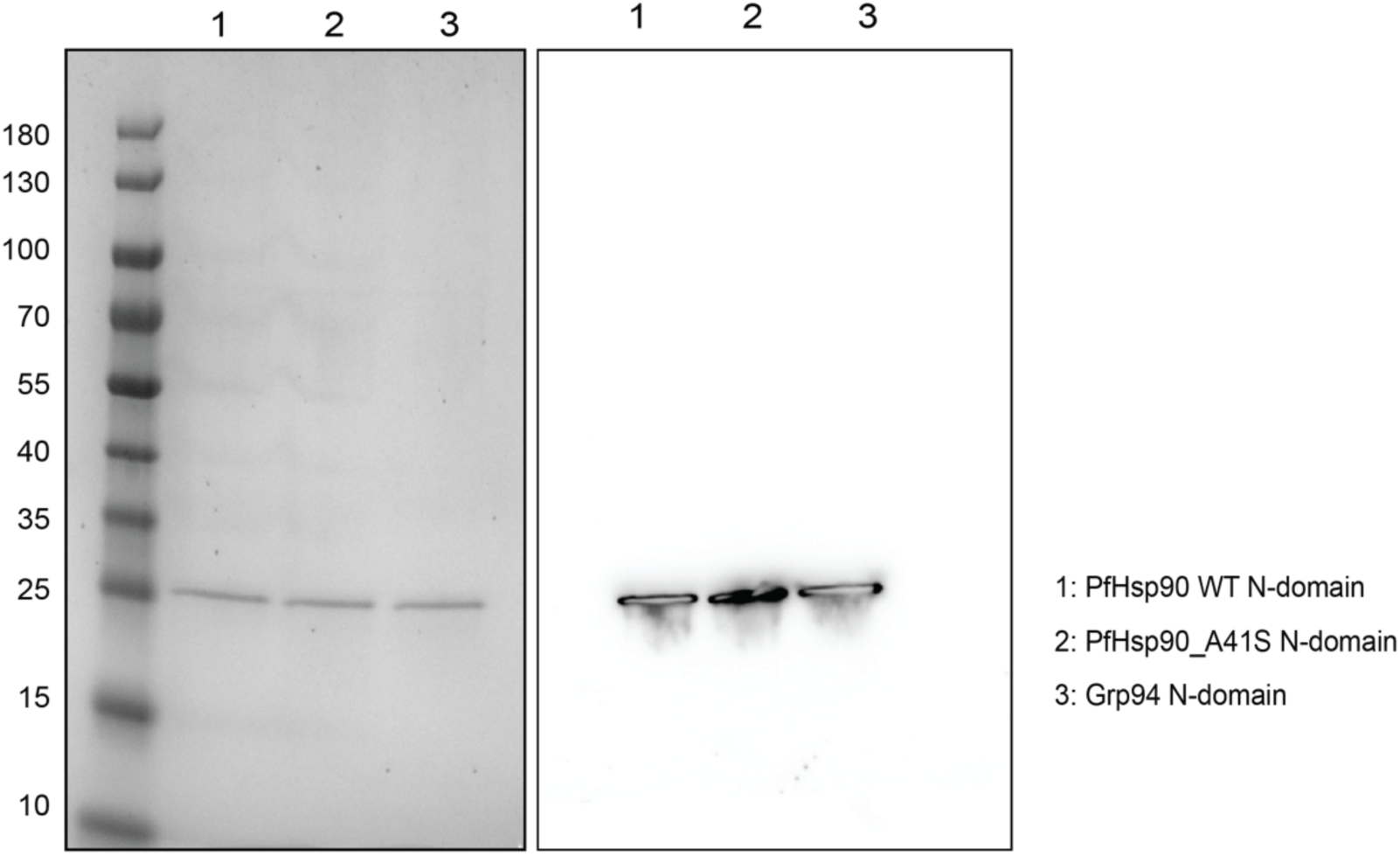
Protein purification validation. SDS-PAGE analysis showing purified proteins. Left: Coomassie blue staining. Right: Anti-His Western blot confirming successful purification of PfHsp90 wild-type N-domain, PfHSP90 A41S N-domain, and PfGRP94 N-domain.

**Figure S6.**
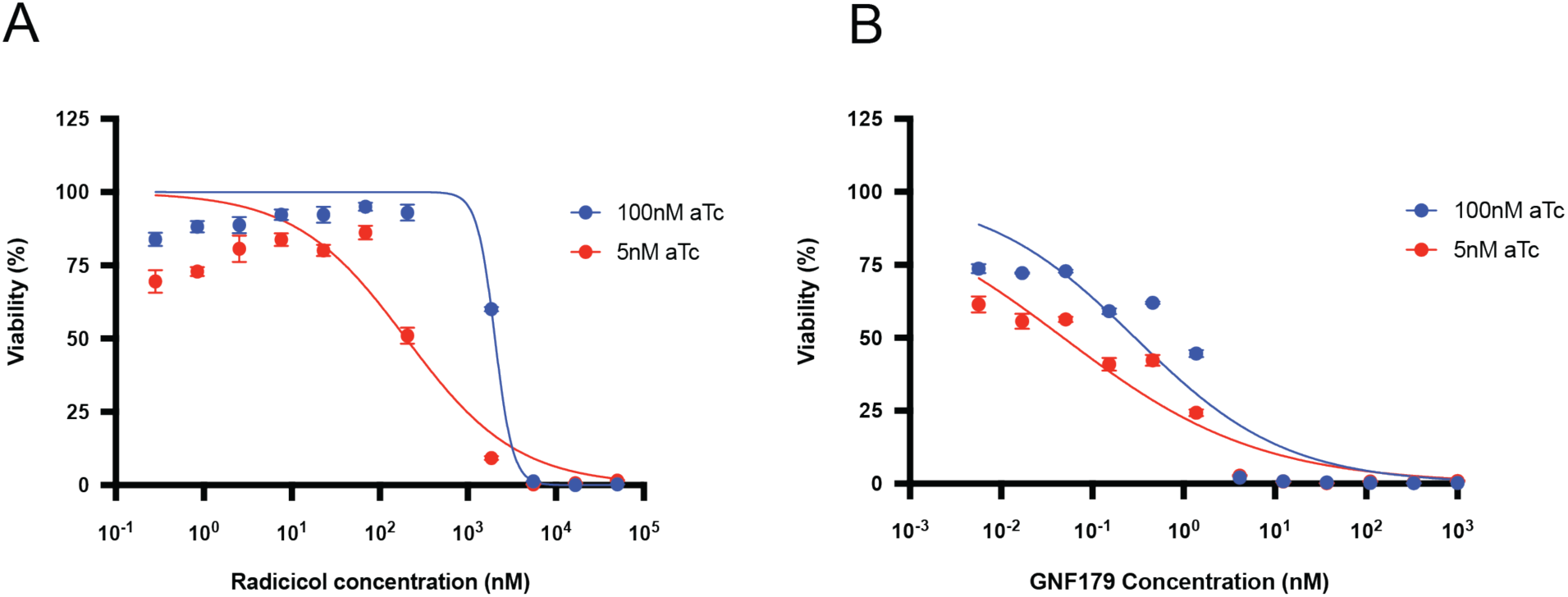
HSP90 conditional knockdown effects on radicicol and GNF179 sensitivity. **(A)** IC₅₀ shift assay for radicicol under HSP90 knockdown conditions comparing 5 nM vs 100 nM aTc treatment. (B) IC₅₀ shift assay for GNF179 under HSP90 knockdown conditions comparing 5 nM vs 100 nM aTc treatment.

## Reference

Adjalley, Sophie, and Marcus Chee San Lee. 2022. “CRISPR/Cas9 Editing of the Plasmodium Falciparum Genome.” In Malaria Immunology, edited by Anja Tatiana Ramstedt Jensen and Lars Hviid, vol. 2470. Methods in Molecular Biology. Springer US. 10.1007/978-1-0716-2189-9_17.

Bridgford, Jessica L., Stanley C. Xie, Simon A. Cobbold, et al. 2018. “Artemisinin Kills Malaria Parasites by Damaging Proteins and Inhibiting the Proteasome.” Nature Communications 9 (1): 3801. 10.1038/s41467-018-06221-1.

Carrasquilla, Manuela, Ndey F. Drammeh, Mukul Rawat, et al. 2022. “Barcoding Genetically Distinct Plasmodium Falciparum Strains for Comparative Assessment of Fitness and Antimalarial Drug Resistance.” mBio 13 (5): e00937–22. 10.1128/mbio.00937-22.

Corbett, Kevin D., and James M. Berger. 2010. “Structure of the ATP-binding Domain of *Plasmodium Falciparum* Hsp90.” *Proteins: Structure*, Function, and Bioinformatics 78 (13): 2738–44. 10.1002/prot.22799.

De Lera Ruiz, Manuel, Paola Favuzza, Zhuyan Guo, et al. 2022. “The Invention of WM382, a Highly Potent PMIX/X Dual Inhibitor toward the Treatment of Malaria.” ACS Medicinal Chemistry Letters 13 (11): 1745–54. 10.1021/acsmedchemlett.2c00355.

Deni, Ioanna, Barbara H. Stokes, Kurt E. Ward, et al. 2023. “Mitigating the Risk of Antimalarial Resistance via Covalent Dual-Subunit Inhibition of the Plasmodium Proteasome.” Cell Chemical Biology 30 (5): 470–485.e6. 10.1016/j.chembiol.2023.03.002.

DiCarlo, James E., Julie E. Norville, Prashant Mali, Xavier Rios, John Aach, and George M. Church. 2013. “Genome Engineering in Saccharomyces Cerevisiae Using CRISPR-Cas Systems.” Nucleic Acids Research 41 (7): 4336–43. 10.1093/nar/gkt135.

Diedrich, Konrad, Bennet Krause, Ole Berg, and Matthias Rarey. 2023. “PoseEdit: Enhanced Ligand Binding Mode Communication by Interactive 2D Diagrams.” Journal of Computer-Aided Molecular Design 37 (10): 491–503. 10.1007/s10822-023-00522-4.

Duffey, Maëlle, Benjamin Blasco, Jeremy N. Burrows, Timothy N.C. Wells, David A. Fidock, and Didier Leroy. 2021. “Assessing Risks of Plasmodium Falciparum Resistance to Select Next-Generation Antimalarials.” Trends in Parasitology 37 (8): 709–21. 10.1016/j.pt.2021.04.006.

Felip, Enriqueta, Fabrice Barlesi, Benjamin Besse, et al. 2018. “Phase 2 Study of the HSP-90 Inhibitor AUY922 in Previously Treated and Molecularly Defined Patients with Advanced Non–Small Cell Lung Cancer.” Journal of Thoracic Oncology 13 (4): 576–84. 10.1016/j.jtho.2017.11.131.

Friesner, Richard A., Jay L. Banks, Robert B. Murphy, et al. 2004. “Glide: A New Approach for Rapid, Accurate Docking and Scoring. 1. Method and Assessment of Docking Accuracy.” Journal of Medicinal Chemistry 47 (7): 1739–49. 10.1021/jm0306430.

Fu, Ying, Leann Tilley, Shannon Kenny, and Nectarios Klonis. 2010. “Dual Labeling with a Far Red Probe Permits Analysis of Growth and Oxidative Stress in *P. Falciparum* -infected Erythrocytes.” Cytometry Part A 77A (3): 253–63. 10.1002/cyto.a.20856.

Ganesan, Suresh M., Alejandra Falla, Stephen J. Goldfless, Armiyaw S. Nasamu, and Jacquin C. Niles. 2016. “Synthetic RNA–Protein Modules Integrated with Native Translation Mechanisms to Control Gene Expression in Malaria Parasites.” Nature Communications 7 (1): 10727. 10.1038/ncomms10727.

Genheden, Samuel, and Ulf Ryde. 2015. “The MM/PBSA and MM/GBSA Methods to Estimate Ligand-Binding Affinities.” Expert Opinion on Drug Discovery 10 (5): 449–61. 10.1517/17460441.2015.1032936.

Haldar, Kasturi, Souvik Bhattacharjee, and Innocent Safeukui. 2018. “Drug Resistance in Plasmodium.” Nature Reviews Microbiology 16 (3): 156–70. 10.1038/nrmicro.2017.161.

Hekkelman, Maarten L., Ida De Vries, Robbie P. Joosten, and Anastassis Perrakis. 2023. “AlphaFill: Enriching AlphaFold Models with Ligands and Cofactors.” Nature Methods 20 (2): 205–13. 10.1038/s41592-022-01685-y.

Janes, Jeff, Megan E. Young, Emily Chen, et al. 2018. “The ReFRAME Library as a Comprehensive Drug Repurposing Library and Its Application to the Treatment of Cryptosporidiosis.” Proceedings of the National Academy of Sciences 115 (42): 10750–55. 10.1073/pnas.1810137115.

Kim, Joungnam, Sara Felts, Laura Llauger, et al. 2004. “Development of a Fluorescence Polarization Assay for the Molecular Chaperone Hsp90.” SLAS Discovery 9 (5): 375–81. 10.1177/1087057104265995.

LaMonte, Gregory M., Frances Rocamora, Danushka S. Marapana, et al. 2020. “Pan-Active Imidazolopiperazine Antimalarials Target the Plasmodium Falciparum Intracellular Secretory Pathway.” Nature Communications 11 (1): 1780. 10.1038/s41467-020-15440-4.

Lawong, Aloysus, Suraksha Gahalawat, Sneha Ray, et al. 2024. “Identification of Potent and Reversible Piperidine Carboxamides That Are Species-Selective Orally Active Proteasome Inhibitors to Treat Malaria.” Cell Chemical Biology 31 (8): 1503–1517.e19. 10.1016/j.chembiol.2024.07.001.

Li, Yang, Lihua Zou, Qiyuan Li, et al. 2010. “Amplification of LAPTM4B and YWHAZ Contributes to Chemotherapy Resistance and Recurrence of Breast Cancer.” Nature Medicine 16 (2): 214–18. 10.1038/nm.2090.

Li, Zi-Nan, and Ying Luo. 2023. “HSP90 Inhibitors and Cancer: Prospects for Use in Targeted Therapies (Review).” Oncology Reports 49 (1): 6. 10.3892/or.2022.8443.

Lim, Michelle Yi-Xiu, Gregory LaMonte, Marcus C. S. Lee, et al. 2016. “UDP-Galactose and Acetyl-CoA Transporters as Plasmodium Multidrug Resistance Genes.” Nature Microbiology 1 (12): 16166. 10.1038/nmicrobiol.2016.166.

Love, Michael I., Wolfgang Huber, and Simon Anders. 2014. “Moderated Estimation of Fold Change and Dispersion for RNA-Seq Data with DESeq2.” Genome Biology 15 (12): 550. 10.1186/s13059-014-0550-8.

Makhoba, Xolani H., Claudio Viegas, Rebamang A. Mosa, Flávia P. D. Viegas, and Ofentse J. Pooe. 2020. “Potential Impact of the Multi-Target Drug Approach in the Treatment of Some Complex Diseases.” *Drug Design*, Development and Therapy 14: 3235–49. 10.2147/DDDT.S257494.

Mansfield, Christopher R., Baiyi Quan, Michael E. Chirgwin, et al. 2024. “Selective Targeting of Plasmodium Falciparum Hsp90 Disrupts the 26S Proteasome.” Cell Chemical Biology 31 (4): 729–742.e13. 10.1016/j.chembiol.2024.02.008.

Millson, Stefan H., Chun-Song Chua, S. Mark Roe, et al. 2011. “Features of the Streptomyces Hygroscopicus HtpG Reveal How Partial Geldanamycin Resistance Can Arise with Mutation to the ATP Binding Pocket of a Eukaryotic Hsp90.” FASEB Journal: Official Publication of the Federation of American Societies for Experimental Biology 25 (11): 3828–37. 10.1096/fj.11-188821.

Murillo-Solano, Claribel, Chunmin Dong, Cecilia G. Sanchez, and Juan C. Pizarro. 2017. “Identification and Characterization of the Antiplasmodial Activity of Hsp90 Inhibitors.” Malaria Journal 16 (1): 292. 10.1186/s12936-017-1940-7.

Muzenda, Florence Lisa, Melissa Louise Stofberg, Wendy Mthembu, Ikechukwu Achilonu, Erick Strauss, and Tawanda Zininga. 2025. “Characterization and Inhibition of the Chaperone Function of *Plasmodium Falciparum* Glucose-Regulated Protein 94 KDA ( *Pf* GRP94 ).” Proteins: Structure, Function, and Bioinformatics 93 (5): 957–71. 10.1002/prot.26779.

Nasamu, Armiyaw S., Alejandra Falla, Charisse Flerida A. Pasaje, et al. 2021. “An Integrated Platform for Genome Engineering and Gene Expression Perturbation in Plasmodium Falciparum.” Scientific Reports 11 (1): 342. 10.1038/s41598-020-77644-4.

Ottilie, Sabine, Madeline R. Luth, Erich Hellemann, et al. 2022. “Adaptive Laboratory Evolution in S. Cerevisiae Highlights Role of Transcription Factors in Fungal Xenobiotic Resistance.” Communications Biology 5 (1): 128. 10.1038/s42003-022-03076-7.

Özen, Ayşegül, and Celia A. Schiffer. 2017. “Substrate-Envelope-Guided Design of Drugs with a High Barrier to the Evolution of Resistance.” In Handbook of Antimicrobial Resistance, edited by Albert Berghuis, Greg Matlashewski, Mark A. Wainberg, and Donald Sheppard. Springer New York. 10.1007/978-1-4939-0694-9_9.

Phillips, Margaret A., Julie Lotharius, Kennan Marsh, et al. 2015. “A Long-Duration Dihydroorotate Dehydrogenase Inhibitor (DSM265) for Prevention and Treatment of Malaria.” Science Translational Medicine 7 (296): 296ra111. 10.1126/scitranslmed.aaa6645.

Posfai, Dora, Amber L. Eubanks, Allison I. Keim, et al. 2018. “Identification of Hsp90 Inhibitors with Anti-Plasmodium Activity.” Antimicrobial Agents and Chemotherapy 62 (4): e01799–17. 10.1128/AAC.01799-17.

Prodromou, C., S. M. Roe, R. O’Brien, J. E. Ladbury, P. W. Piper, and L. H. Pearl. 1997. “Identification and Structural Characterization of the ATP/ADP-Binding Site in the Hsp90 Molecular Chaperone.” Cell 90 (1): 65–75. 10.1016/s0092-8674(00)80314-1.

Ramdhave, Anup S., Dhaval Patel, I. Ramya, Mukesh Nandave, and Prashant S. Kharkar. 2013. “Targeting Heat Shock Protein 90 for Malaria.” Mini Reviews in Medicinal Chemistry 13 (13): 1903–20. 10.2174/13895575113136660094.

Rosenthal, Philip J., Victor Asua, and Melissa D. Conrad. 2024. “Emergence, Transmission Dynamics and Mechanisms of Artemisinin Partial Resistance in Malaria Parasites in Africa.” Nature Reviews Microbiology 22 (6): 373–84. 10.1038/s41579-024-01008-2.

Roy, Nainita, Rishi Kumar Nageshan, Shatakshi Ranade, and Utpal Tatu. 2012. “Heat Shock Protein 90 from Neglected Protozoan Parasites.” Biochimica Et Biophysica Acta 1823 (3): 707–11. 10.1016/j.bbamcr.2011.12.003.

Siqueira-Neto, Jair L., Kathryn J. Wicht, Kelly Chibale, Jeremy N. Burrows, David A. Fidock, and Elizabeth A. Winzeler. 2023. “Antimalarial Drug Discovery: Progress and Approaches.” Nature Reviews Drug Discovery 22 (10): 807–26. 10.1038/s41573-023-00772-9.

Southworth, Daniel R., and David A. Agard. 2008. “Species-Dependent Ensembles of Conserved Conformational States Define the Hsp90 Chaperone ATPase Cycle.” Molecular Cell 32 (5): 631–40. 10.1016/j.molcel.2008.10.024.

Stofberg, Melissa Louise, Florence Lisa Muzenda, Ikechukwu Achilonu, Erick Strauss, and Tawanda Zininga. 2024. “*In Silico* Screening of Selective ATP Mimicking Inhibitors Targeting the *Plasmodium Falciparum* Grp94.” *Journal of Biomolecular Structure and Dynamics*, March 18, 1–12. 10.1080/07391102.2024.2329304.

Suzuki, Yo, Robert P St Onge, Ramamurthy Mani, et al. 2011. “Knocking out Multigene Redundancies via Cycles of Sexual Assortment and Fluorescence Selection.” Nature Methods 8 (2): 159–64. 10.1038/nmeth.1550.

Swann, Justine, Victoria Corey, Christina A. Scherer, et al. 2016. “High-Throughput Luciferase-Based Assay for the Discovery of Therapeutics That Prevent Malaria.” ACS Infectious Diseases 2 (4): 281–93. 10.1021/acsinfecdis.5b00143.

Tye, Mark A., N. Connor Payne, Catrine Johansson, et al. 2022. “Elucidating the Path to Plasmodium Prolyl-tRNA Synthetase Inhibitors That Overcome Halofuginone Resistance.” Nature Communications 13 (1): 4976. 10.1038/s41467-022-32630-4.

Venkatesan, Priya. 2025. “WHO World Malaria Report 2024.” The Lancet Microbe 6 (4): 101073. 10.1016/j.lanmic.2025.101073.

Whitesell, Luke, Nicole Robbins, David S. Huang, et al. 2019. “Structural Basis for Species-Selective Targeting of Hsp90 in a Pathogenic Fungus.” Nature Communications 10 (1): 402. 10.1038/s41467-018-08248-w.

XX. Schrödinger Release 2024-3: Maestro, Schrödinger, LLC, New York, NY, 2024.

